# Investigation of betaine as a novel psychotherapeutic for schizophrenia

**DOI:** 10.1101/614164

**Authors:** Tetsuo Ohnishi, Shabeesh Balan, Manabu Toyoshima, Motoko Maekawa, Hisako Ohba, Akiko Watanabe, Yoshimi Iwayama, Chie Shimamoto-Mitsuyama, Yayoi Nozaki, Yasuko Hisano, Kayoko Esaki, Atsuko Nagaoka, Junya Matsumoto, Mizuki Hino, Nobuko Mataga, Akiko Hayashi-Takagi, Yasuto Kunii, Akiyoshi Kakita, Hirooki Yabe, Takeo Yoshikawa

## Abstract

Betaine is known to act against various biological stresses and its levels were reported to be decreased in schizophrenia patients. Using *Chdh* (a gene for betaine synthesis)-deficient mice and betaine-supplemented inbred mice, we assessed the role of betaine in psychiatric pathophysiology, and its potential as a novel psychotherapeutic, by leveraging metabolomics, behavioral-, transcriptomics and DNA methylation analyses. The *Chdh*-deficient mice revealed remnants of psychiatric behaviors along with schizophrenia-related molecular perturbations. Betaine supplementation elicited genetic background-dependent improvement in cognitive performance, and suppressed methamphetamine (MAP)-induced behavioral sensitization. Furthermore, betaine rectified the altered antioxidative and proinflammatory responses induced by MAP and *in vitro* phencyclidine treatments. Notably, betaine levels were decreased in the postmortem brains from schizophrenia, and a coexisting elevated carbonyl stress, a form of oxidative stress, demarcated a subset of schizophrenia with “betaine deficit-oxidative stress pathology”. We revealed the decrease of betaine levels in glyoxylase 1 (*GLO1*)-defícient hiPSCs, which shows elevated carbonyl stress, and the efficacy of betaine in alleviating it, thus supporting a causal link between betaine and oxidative stress conditions. Furthermore, a *CHDH* variant, rs35518479, was identified as a cis-expression quantitative trait locus (QTL) for *CHDH* expression in postmortem brains from schizophrenia, allowing genotype-based stratification of schizophrenia patients for betaine efficacy. In conclusion, the present study underscores the potential benefit of betaine in a subset of schizophrenia.

## Introduction

There is an urgent need for novel medicines with new actions for psychiatric illness, because almost all of the present therapeutics are designed to act on the monoamine receptors or transporters (1), and there is a substantial population where the presently available drugs are unsatisfactory (2). Altered brain metabolome and its reflection in peripheral samples is a growing observation in schizophrenia, which could be beneficial for deciphering molecular pathophysiology of schizophrenia and thereby identifying novel druggable targets (3–5). Betaine (glycine betaine or trimethylglycine) is a metabolite that is reported to be decreased in the plasma sample of patients with first-episode schizophrenia (6), thus tempting the examination of its putative role in psychiatric conditions.

In vertebrates, betaine is ingested from the diet (5) and is also endogenously synthesized in mitochondria from its precursor choline using choline dehydrogenase (CHDH) (Fig. 1) (4). To date, several biological functions of betaine have been proposed, which include (i) osmotic regulator (compatible solute) (7, 8); (ii) antioxidative/anti-inflammatory activity (9); (iii) supplier of methyl donor *S*-adenosylmethionine (10); and (iv) mitigator of noxious elevated homocysteine (11) (Supplementary Fig. 1). The latter two are endowed through acceleration of the turnover of the methionine–homocysteine cycle (constituting one-carbon metabolism together with the folate cycle), where betaine serves as a substrate in the betaine–homocysteine *S*-methyltransferase (BHMT) reaction, converting homocysteine to methionine (12) (Fig. 1).

**Figure 1.**
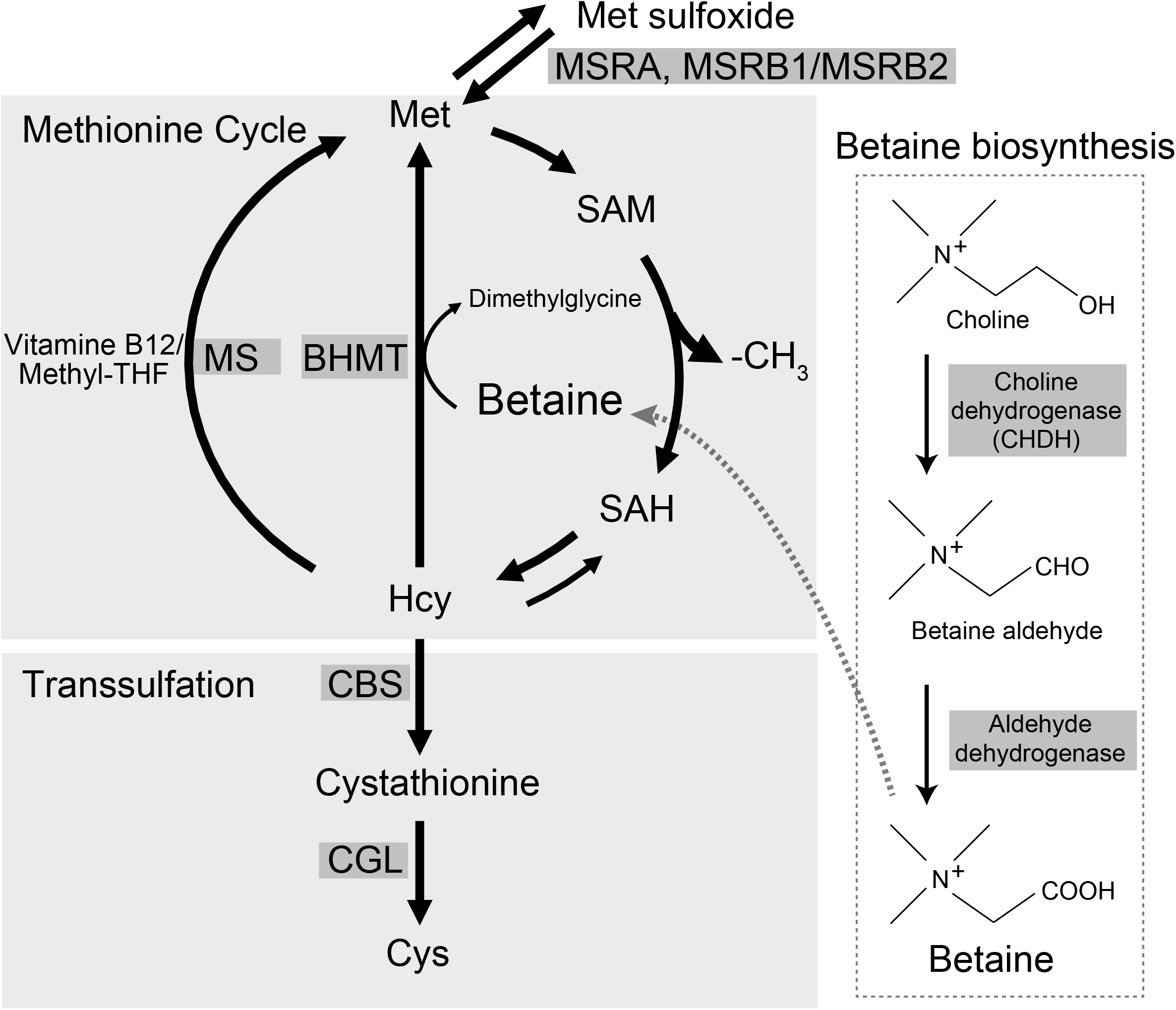
Methionine–homocysteine cycle and biosynthesis of betaine. The methionine–homocysteine cycle, betaine biosynthesis pathway, and transsulfation pathway are schematically presented. Met, methionine; SAM, *S*-adenosylmethionine; Met sulfoxide, methionine sulfoxide; -CH_3_, methyl group; SAH, *S*-adenosylhomocysteine; Hcy, homocysteine; methyl-THF, 5-methyltetrahydrofolate; MS, methionine synthase; CBS, cystathionine beta-synthase; CGL, cystathionine beta-lyase; MSRA, methionine sulfoxide reductase A; MSRB1/2, methionine sulfoxide reductase B1/2.

In this study, we aimed to test the role of betaine in schizophrenia pathophysiology, and to evaluate its potential as a novel psychotherapeutic. Using a *Chdh* (a gene for betaine synthesis)-deficient mice and betaine-supplemented inbred mice, we assessed betaine-related metabolite dynamics and the effects of betaine deficits and supplementation, by leveraging metabolomics, behavioral-, transcriptomics and DNA methylation analyses. Further, we queried whether betaine deficits and the associated pathology are observed in postmortem brains from schizophrenia. We examined the efficacy of betaine in human induced pluripotent stem cells (hiPSCs) manifesting elevated carbonyl stress, which is a tributary of oxidative stress and caused by accumulation of “advanced glycation end products” (AGEs) (13) (Supplementary Fig. 2). The enhanced carbonyl stress is known to underlie the pathophysiology of a subset of schizophrenia (14, 15). Lastly, we searched for genetic variants that can be used as a biomarker to predict “betaine-responders”.

## Methods

See the Supplementary information for the details of the techniques outlined below.

### Study approval

All the animal experiments were performed in compliance with relevant laws, and guidelines were approved by the Animal Ethics Committee at RIKEN Center for Brain Science, Japan (H29-2-204(3) and 2016-058(4)). Human induced pluripotent stem cell study was approved by the Human Ethics Committee at RIKEN for iPSC study (Wako-daisan 25-14). Experiments in postmortem brain samples were approved by Fukushima Medical University, Japan (1685 and 2381) and Niigata University School of Medicine, Japan (G2015-0827).

### Animals

The inbred C57BL/6NCrl (B6N) and C3H/HeNCrl (C3HN) mouse strains were obtained from Japan’s Charles River Laboratories (Yokohama, Japan). The animals were housed in groups of four in standard cages, in a temperature and humidity-controlled room with a 12 h light/dark cycle (lights on at 08:00). The animals had free access to standard lab chow and tap water. All animal experiments were done using male animals (six to 15 animals/group, depending on experiments) between 9:30 am and 5:00 pm. In the study using the *Chdh* KO mice, the wild-type littermates by intercross between heterozygotes were used as control. All the analyses were conducted between nine and 14 weeks of age.

### Postmortem Brain Samples

Postmortem brain tissues (BA17; Brodmann Area 17) from schizophrenia and age-matched control samples were obtained from the Postmortem Brain Bank of Fukushima for Psychiatric Research and Brain Research Institute, Niigata University, Japan (in total *n* = 24 for schizophrenia and *n* = 31 for control) (16–18). Each individual with schizophrenia fulfilled the diagnostic criteria established by the American Psychiatric Association (Diagnostic and Statistical Manual of Mental Disorders: DSM-IV) and had no history of any other neurological disorder or substance abuse. The age matched control samples were with no history of neuropsychiatric disorders. Also, neuropathological evaluation in the control brain samples ruled out any pathological abnormalities characteristic of neurological disorders, though some brains showed indication of mild senility.

### Generation of *Chdh*-deficient Mice

The *Chdh*-deficient mice were generated by the genome editing methodology using the CRISPR/Cas9 nickase.

### Estimation of Betaine and Other Metabolites

Choline, methionine, betaine, cystathionine, cystine, *S*-adenosylmethionine (SAM), and *S*-adenosylhomocysteine (SAH) levels were estimated by liquid chromatography-mass spectrometry (LC/MS). Cysteine, homocysteine and glutathione (GSH) were measured as their reduced forms by high performance liquid chromatography (HPLC). The plasma levels of blood urea nitrogen (BUN), creatinine (CRE), aspartate transaminase (AST), alanine transaminase (ALT), lactate dehydrogenase (LDH) and cholinesterase (CHE) were measured using dry-chemistry analysis (Fuji Dri-Chem 3500V; FUJIFILM Medical Co., Ltd., Tokyo, Japan).

### Behavioral Analyses

All behavioral tests relevant to psychiatric illnesses, except for the methamphetamine (MAP)-induced behavioral sensitization test, were performed according to the previously published methods (19). For the MAP-induced behavioral sensitization test, see the Supplementary Methods.

### DNA Methylation Analysis

DNA methylation in frontal cortex of *Chdh* knockout (KO) mice (*n* = 6) and wild-type (WT) control (*n* = 6) was performed by targeted methylation sequencing of 109 Mb of mouse genomic regions, which included CpG islands, known tissue-specific differentially methylated regions (DMR), open regulatory annotations and Ensembl regulatory features.

### RNA-seq Analysis

Transcriptome analysis in frontal cortical brain region was performed by RNA-seq in (a) *Chdh* KO mice (*n* = 6) versus WT controls (*n* = six), and (b) B6N and C3HN mouse strains administered betaine in comparison to respective controls administered water (*n* = 6 in each group).

### Real-time Quantitative Reverse Transcription (RT)-PCR

Targeted gene expression was measured by real-time quantitative RT-PCR using TaqMan assays (20).

### Rat Primary Cortical Neuron Culture

Rat primary cortical neuron was isolated from the cortices of Sprague-Dawley rats (obtained from Japan’s Charles River) at embryonic day 18.5 (E18.5) as described previously (21). For the details, see the Supplementary Methods.

### Establishment of *GLO1*-deficient hiPSCs

Human induced pluripotent stem cells (hiPSCs) were established from peripheral blood mononuclear cells using Sendai virus vector (22). Isogenic *GLO1*-deficient hiPSCs were generated using CRISPR/Cas9-mediated genome editing.

### Carbonyl Stress Assay

Carbonyl Stress was analyzed by Western blotting using anti-AGE antibody. See the Supplementary Methods.

### *Cis*-eQTL analysis

For eQTL analysis, we selected *BHMT1, CHDH* and *GLO1* and downloaded eQTL data for the brain tissues from GTEx portal (https://gtexportal.org/home/) (Release V7; dbGaP Accession phs000424.v7.p2). The significant eQTL variants with high effect size were selected and filtered for minor allele frequency (MAF) > 30% in Japanese population with reference to the Tohuku megabank genome variation data from 3,552 whole genome sequences (https://jmorp.megabank.tohoku.ac.jp/201902/downloads) (3.5KJPNv2). These variants were further pruned for linkage disequilibrium (LD) status (*r*^2^ < 0.4), yielding variants from independent LD blocks. The shortlisted variants (4 in *BHMT*, 3 in *CHDH* and 3 in *GLO1*) were genotyped in schizophrenia postmortem brain tissues (BA17) (*n* = 50) by TaqMan SNP genotyping Assays (23). Gene expression was measured by real-time quantitative RT-PCR using TaqMan assays. Outliers (more or less than mean ± 2SD) were excluded. Association of the variants with the gene expression and metabolites were tested by one-way ANOVA and Student’s *t*-test.

### Statistics

All values in the figures represent the mean ± SEM. Statistical analysis and graphical representation were performed using GraphPad Prism 6 (GraphPad Software). The total sample size (*n*) was described in the respective figure legends. Statistical significance was determined using a two-tailed Student’s *t*-test. When multiple comparisons were needed, one-way or two-way ANOVA with Fisher’s least significant difference (LSD) test, Tukey’s multiple comparison test, Bonferroni’s correction, or Dunnett’s multiple comparison test was used as indicated in the figure legends. A *P* value of less than 0.05 was considered as statistically significant.

## Results

### Generation of *Chdh*-deficient mice

In vertebrates, betaine is biosynthesized by two biochemical reactions (Fig. 1). The first step, oxidation of choline to betaine aldehyde, is mediated by choline dehydrogenase encoded by the *Chdh*/*CHDH* gene, and in turn, betaine aldehyde is converted into betaine by betaine aldehyde dehydrogenase. To create an animal model with lowered betaine level, the first coding exon, exon 2 of the *Chdh* gene, was targeted by the CRISPR/Cas9n on the genetic background of the inbred B6N mouse (Supplementary Fig. 3A and B). Homozygotes for the gene disruption seemed healthy and revealed no clear abnormalities in growth or morphology (Supplementary Fig. 3C).

### Betaine levels and effects of betaine supplementation in *Chdh*-deficient mice

The disruption of the *Chdh* gene significantly reduced the betaine levels in the frontal cortex from ~20 pmol/mg tissue to levels near detection limit (*p* < 0.01) (Fig. 2A). This suggested that the endogenous biosynthesis rather than intake from diet maintained the level of betaine in the brain. Supplementation of betaine via drinking water (2.17% betaine, 2.5% as betaine monohydrate) for 3 weeks, significantly increased betaine level in the brain of both the WT (*p* < 0.0001) and gene-disrupted mice (*p* < 0.001) (Fig. 2A), indicating that peripherally-administered betaine can penetrate the blood–brain barrier and enter the brain.

**Figure 2.**
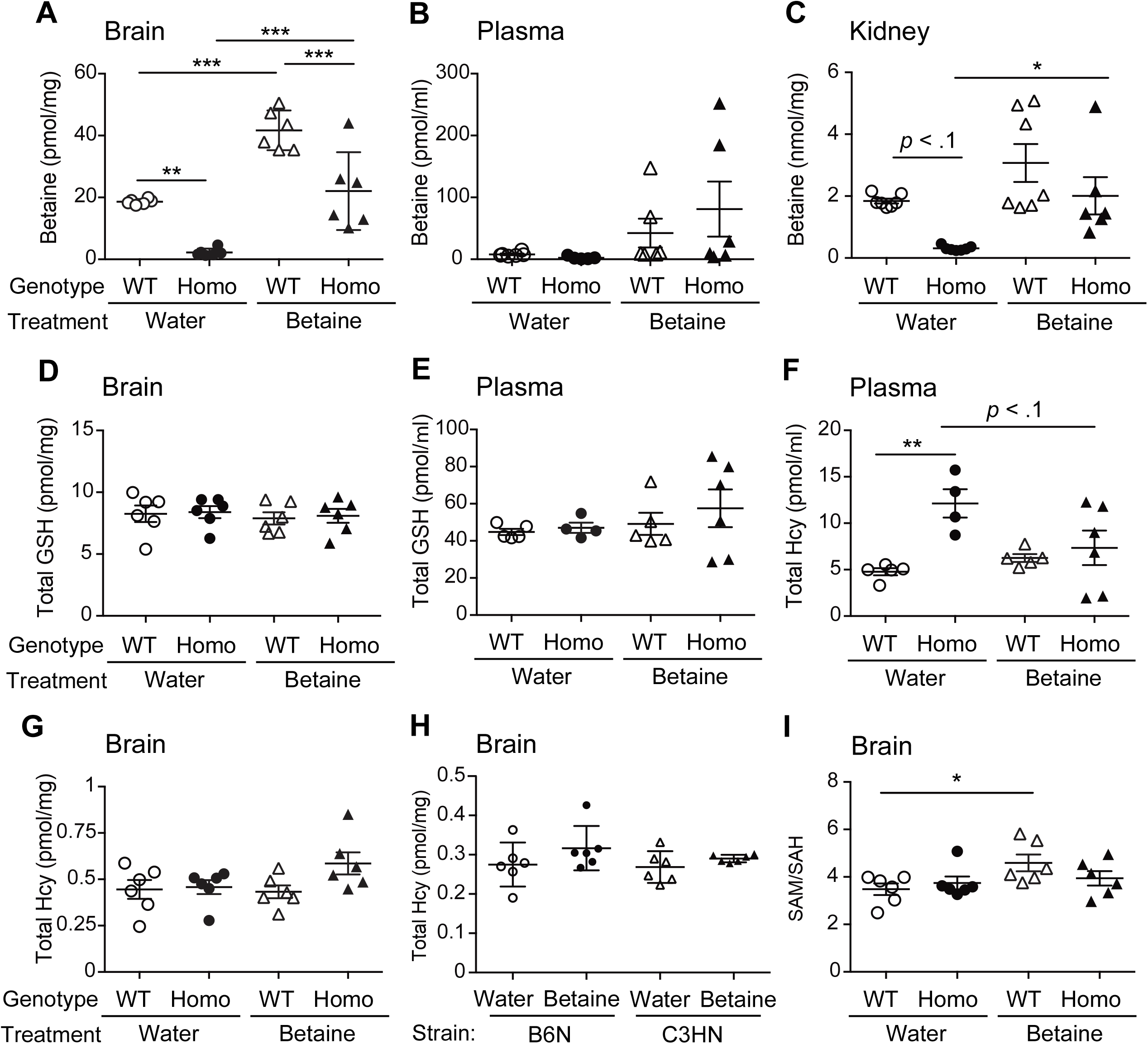
Effects of *Chdh*-deficiency and betaine supplementation on the metabolite contents. Metabolite contents were measured in the *Chdh*-deficient (Homo) and wild-type *Chdh* littermate control (WT) mice with (betaine) or without (water) chronic betaine supplementation. **(A)** Betaine levels were significantly reduced in the frontal cortex of *Chdh*-deficient mice, which were restored by betaine supplementation. **(B)** In plasma, betaine levels were not statistically significant among the tested groups. **(C)** Betaine levels were drastically reduced in the kidney of *Chdh*-deficient mice and were restored to WT level. Total glutathione (GSH) contents (GSH + GSSG), in the **(D)** brain and **(E)** plasma did not reveal any statistically significant differences among the tested groups. **(F)** Total plasma homocysteine increased in the *Chdh*-deficient mice when compared to WT, and showed a trend toward WT level after betaine supplementation; however, in the brain, the levels of total homocysteine were unchanged in the tested groups **(G-H)**. **(I)** SAM/SAH ratio, indicating the cellular methylation potential, was significantly increased with betaine supplementation in WT B6N mice. Data represent mean ± SEM. \**p* < .05, \*\**p* < .01, ***, *p* < .001; Tukey’s multiple comparison test among the four groups. *n* = five to seven per group.

In the plasma, however, betaine levels were near detection limit in both the WT and *Chdh*-deficient mice in our measurement system (Fig. 2B). Median betaine concentration in plasma in humans has been reported to be 27.8 pmol/L (24), which was in the detectable range in our system. Therefore, the peripheral betaine turnover could be different between mice and humans. Although the plasma betaine levels were increased after the betaine supplementation, the increments were not significant in both the WT and KO animals (Fig. 2B), partially due to widely variable betaine levels in the betaine-administered animals.

### No adverse effects of betaine deficiency in kidney and liver functions

Since the kidney and liver show high expression of Chdh and contain abundant levels of betaine (25), we examined the effects of gene disruption with respect to their functions. The gene disruption elicited a trend of decreased betaine levels (*p* < 0.1) (Fig. 2C). Betaine levels in the kidney of KO mice were replenished to those of WT littermates upon betaine supplementation (Fig. 2C). Betaine in the liver plays a crucial role in one-carbon metabolism (25), but no impairments of liver function were observed by the *Chdh* disruption, as shown by the unaltered plasma levels of AST, ALT, LDH, CHE, BUN or CRE (Supplementary Table 1). Furthermore, the total levels of glutathione (GSH + GSSG) were not altered by the gene disruption (Fig. 2D and E) in the brain or plasma, suggesting that the lack of Chdh activity itself did not invoke a serious oxidative stress condition in mice.

### Levels of metabolites in methionine–homocysteine cycle in the brain

Among the metabolites in the methionine–homocysteine cycle that were driven by betaine (Fig. 1), levels of homocysteine, a potent oxidant, changed differentially between the plasma and brain. In the plasma, homocysteine concentration was increased by the disruption of *Chdh* (*p* < 0.01), and it showed a trend towards WT level (*p* < 0.1) upon betaine supplementation (Fig. 2F). While in the brain (frontal cortex), homocysteine levels were unchanged among the groups (Fig. 2G). We also examined the two inbred mouse strains, B6N (the same genetic background with the *Chdh*-deficient mouse) and C3HN, each of which manifested two extremities for prepulse inhibition function (B6N, highest; C3HN, lowest), a schizophrenia and other mental disorder-related endophenotype (26). Total homocysteine levels in the brain after betaine supplementation was also unchanged in the two inbred strains (Fig. 2H). The results suggest that homocysteine levels in the brain are robustly maintained constant against perturbation of peripheral betaine levels. Interestingly, the cellular methylation capacity deduced from the SAM/SAH ratio (27, 28), showed a significant increase (*p* < 0.05) upon betaine supplementation in the B6N WT brain (Fig. 2I), suggesting a possibility that externally administered betaine could have an impact on the methylation capacity to some extent.

### DNA methylation and transcriptome analyses in the *Chdh*-deficient mice

To explore the impact of betaine on DNA methylation status, a targeted analysis of methylated genomic regions at single base pair resolution in the frontal cortex from *Chdh*-deficient mice was performed. The results revealed no statistically significant differences in DNA methylation when compared to the WT mice after multiple corrections (*q* value > 0.05; Fig. 3A, Supplementary Table 2). As an exploratory approach, we selected differentially methylated gene promoters, employing a conservative estimate of *p* < 0.05 to test the enrichment of canonical pathways and gene ontologies (Fig. 3B, Supplementary Table 3). Significant enrichment was observed for “tight junction signaling” and “sertoli cell-sertoli cell junction signaling” pathways for hypermethylated gene promoters, the latter could be related to the infertility phenotype of *Chdh*-deficient mice (29). In the case of hypomethylated promoters, cytoskeletal dynamics pathways such as “integrin signaling” and “ephrin receptor signaling” were enriched (Fig. 3B).

**Figure 3.**
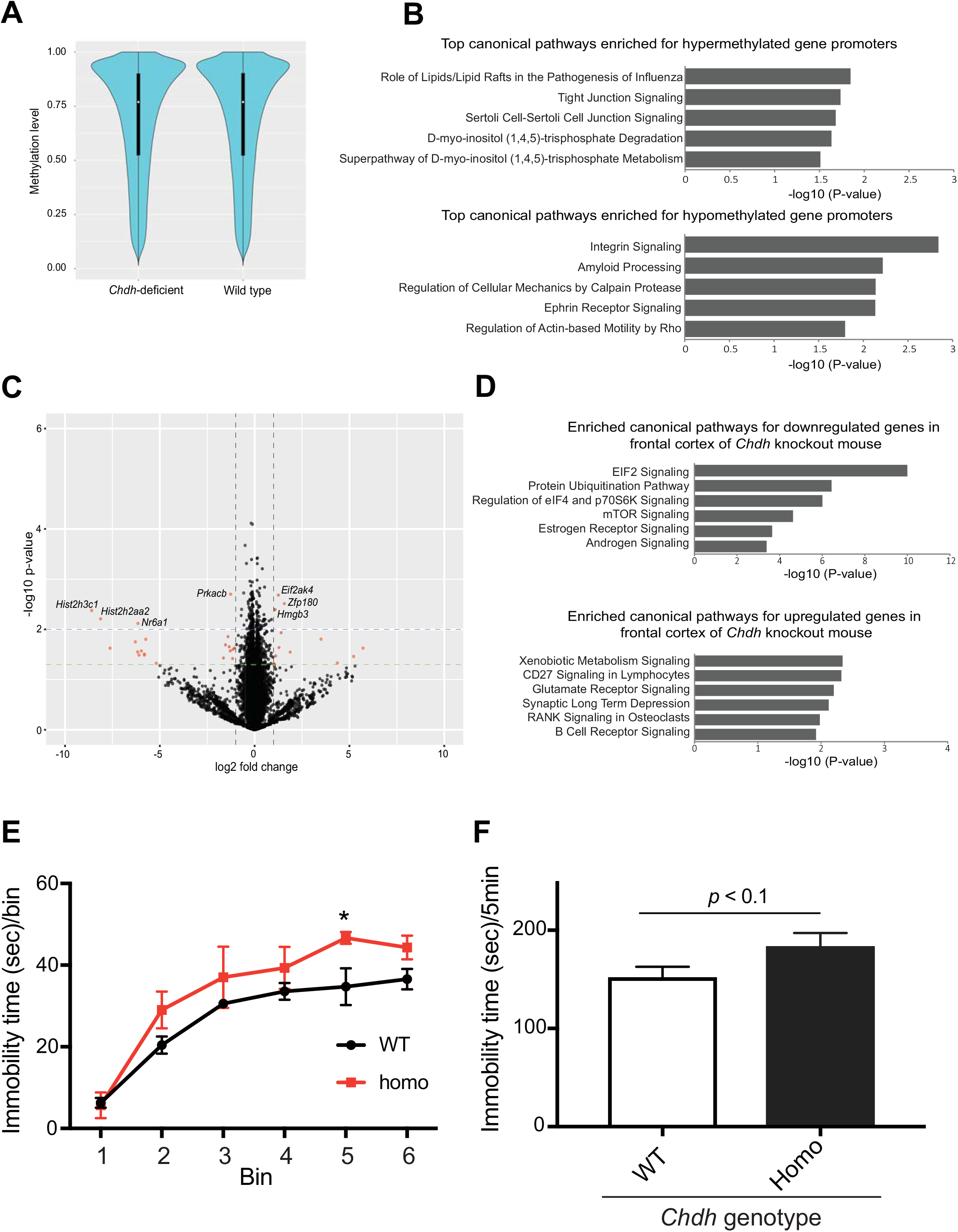
DNA methylome, transcriptome and behavioral analyses of *Chdh*-deficient mice. **(A)** Targeted DNA methylation analysis across the genome, from the frontal cortex of *Chdh*-deficient mice, did not reveal any statistically significant differences (after correcting for multiple tests; *q* value > .05) when compared to the wild-type control. **(B)** Pathway analysis of genes with a *p* < 0.05 (uncorrected) indicated an enrichment of “tight junction signaling” and “sertoli cell-sertoli cell junction signaling” pathways for hypermethylated promoters, implicated in spermatogenesis. Hypomethylated genes were enriched for pathways involving cytoskeletal dynamics; *n* = six in each group. **(C)** Volcano plot shows differentially expressed genes between *Chdh* KO mice and wild type controls from frontal cortex; *n* = six in each group. Green and blue dashed lines indicate *p* value thresholds of 0.05 and 0.01, respectively. Top hits among the differentially expressed genes (*p* < 0.01 and absolute fold-change > 2) are highlighted in the plot. **(D)** Differentially expressed genes (*p* < 0.05) were analyzed for the enriched canonical pathways, which revealed enrichment for signalling pathways involved in the regulation of translational control and protein synthesis/degradation, for downregulated genes. **(E)** Results of forced swim test are presented. 1 bin corresponds to 1 min. Note that the *Chdh*-deficient mice showed increased immobility in bin five (*p* = 0.01). Data represent mean ± SEM. \**p* < 0.05, \*\**p* < 0.01; Fisher’s Least Significant Difference (LSD) test. *n* = 21 per group. **(F)** Cumulative values of immobility time for five minutes (bins two to six) were calculated. Note that KO homozygotes revealed a trend of increase in immobility time (*p* = 0.09). Data represent mean ± SEM. *n* = 21 per group.

Regarding the transcriptomic level changes elicited by the *Chdh* disruption in the frontal cortex, a total of 851 genes were significantly dysregulated (546 upregulated and 305 downregulated) in the KO compared to the WT littermates (*p* < 0.05) (Fig 3C, and Supplementary Table 4). The Ingenuity Pathway Analysis showed significant enrichment for molecular pathways, for example, eukaryotic initiation factor 2 (eIF2) signaling, protein ubiquitination pathway, regulation of eukaryotic initiation factor 4 (eIF4) and p70 S6 kinase signaling, and mammalian target of rapamycin (mTOR) signaling for the downregulated genes (*p* < 0.05) (Fig. 3D, and Supplementary Table 5). These results were in accordance with the reduced protein synthesis observed in schizophrenia, which involve the eIF2α, eIF4 and mTOR signaling pathways (30). Collectively, the altered molecular deficits by the *Chdh* disruption may represent a “precursor signature” (31) underling schizophrenia pathophysiology.

### *Chdh*-deficient mice displayed remnants of depressive behavior

We examined 15 distinct behavioral phenotypes relevant to psychiatric illnesses (Supplementary Fig. 6). In the forced swim test, immobility time at bin 5 (4–5 min) was significantly prolonged (*p* < 0.05), and total immobility time showed a trend of increase (*p* < 0.1) in the *Chdh*-deficient mice compared to the WT littermates (Figs. 3E and F), suggesting a depressive trait in the gene-deficient mice. In the other behavioral tests, no significant differences were observed between the WT and *Chdh*-deficient mice (Supplementary Fig. 6). The limited behavioral deficits elicited by the *Chdh* disruption could be partly due to the fact that betaine was not completely depleted from the whole body, because betaine was available from the diet or through intestinal microflora (32, 33).

### Effects of betaine supplementation in inbred mouse strains

To determine the therapeutic potential of betaine in psychiatric disorders, betaine was administered via drinking water (2.17% betaine) for 3 weeks to the two inbred mouse strains; B6N and C3HN. In general, efficacy of drugs, in particular neurotropics, varies with respect to the individual patients or genetic backgrounds. The same 15 distinct behavioral phenotypes as evaluated in the *Chdh*-deficient mice were measured. Notably, betaine supplementation elicited the differential behavioral effects in the two strains. In the novel object recognition test (NORT), B6N mice showed significantly longer stay time at a novel object in the retention session in the betaine-supplemented group compared to the water-drinking control, suggesting improved cognitive memory performance in the B6N strain (Fig. 4A). Both B6N and C3HN mouse strains showed better performance with betaine supplementation in the Y-maze test, which assessed the spatial working memory ability, compared to the control (Fig. 4B). There were no significant differences between the betaine-administered and control groups in B6N or C3HN animals in the other behavioral tests, and importantly no behavioral measures were impaired by betaine treatments in both strains (data not shown). No gross abnormalities in blood chemistry were also observed (data not shown).

**Figure 4.**
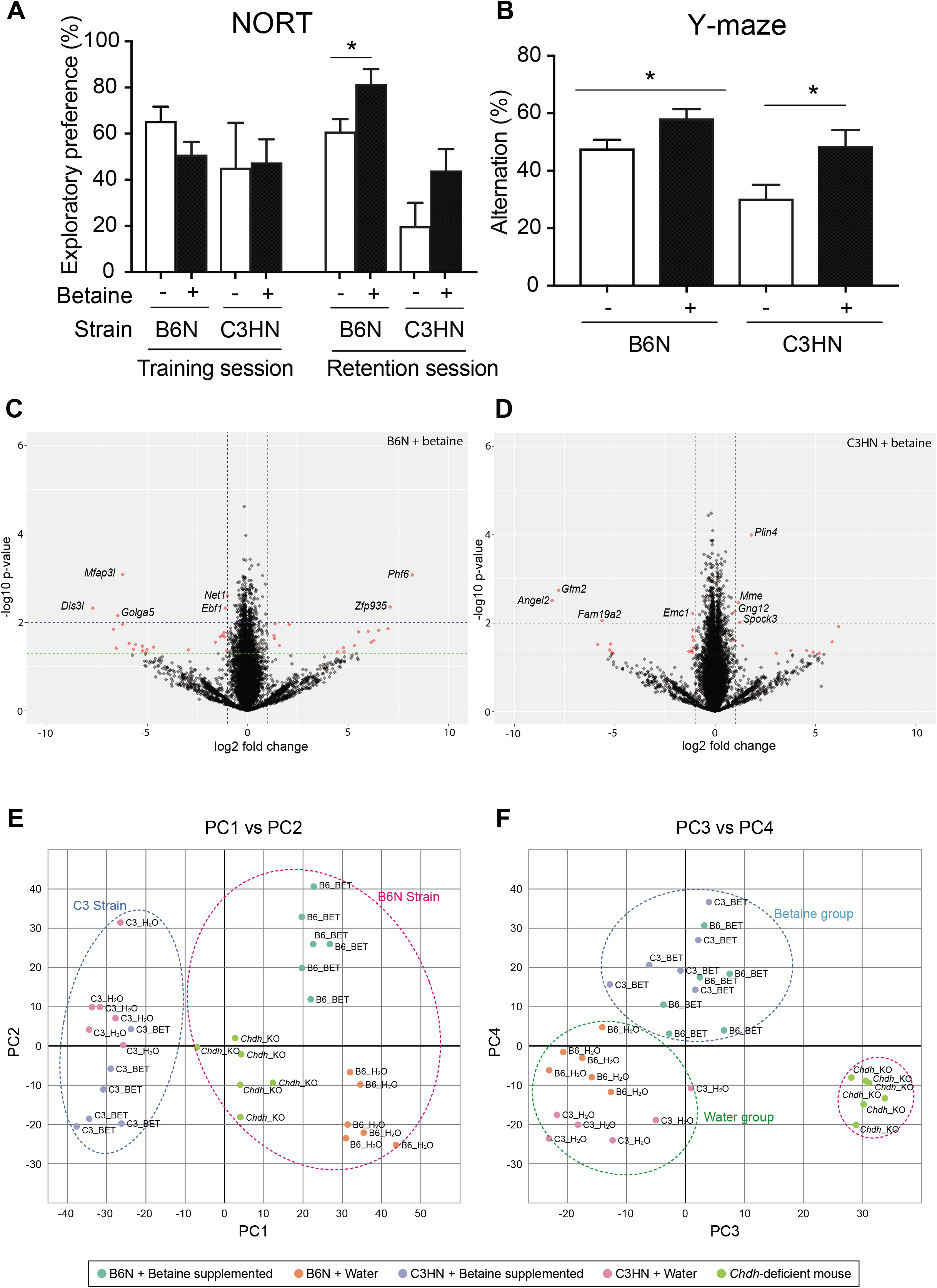
Betaine supplementation improved cognitive performance and elicited gene expressional changes. **(A)** Novel object recognition test (NORT), and (B) Y-maze test were conducted with (+) or without (−) chronic betaine supplementation in two inbred strains, B6N and C3HN. Mouse strains displayed genetic background-dependent improvement in cognitive and spatial memory performance. Data represent mean ± SEM. \**p* < .05; Student’s *t*-test. *n* = seven to ten per group. Volcano plot shows differentially expressed genes from frontal cortex upon betaine supplementation in **(C)** B6N and **(D)** C3HN mice when compared to the controls that were fed water. Green and blue dashed lines indicate *P*-value thresholds of 0.05 and 0.01, respectively. Top hits among the differentially expressed genes (*p* < 0.01 and absolute fold-change > 2) are highlighted in the plot. **(E)** Principal component analysis of transcriptome data (PC 1 vs. 2) shows clear segregation of B6N and C3HN mouse strains, and **(F)** PC 3 vs. 4 shows segregation in relation to betaine supplementation irrespective of strain differences; *n* = six, in each group.

Transcriptome analysis of the frontal cortex revealed that the patterns of gene expression caused by betaine were distinct between the two strains, presumably stemming from the underlying genetic differences and aligning with their behavioral differences (Figs. 4C, D, and Supplementary Table 6). Interestingly, the principal component analysis (PCA) of transcriptome data revealed clear segregation of mouse strains in relation to betaine supplementation (based on principal components 1 and 2) (Fig. 4E). The principal components 3 and 4 could clearly discriminate the effect of betaine supplementation irrespective of strain differences (Fig. 4F). Gene ontology enrichment analysis of genes whose expression was positively correlated with PCA loading factor in these principal components revealed biological processes relevant for the cognition, memory, and neurodevelopment related terms (Supplementary Table 7). Meanwhile, certain common molecular phenotypes were also evident (Supplementary Fig. 7, Supplementary Table 8, and 9). These results highlight an interaction of genetic background × drug (betaine). Gene ontology analysis of upregulated genes by betaine supplementation in both strains identified mitogen-activated protein kinase (MAPK) signaling cascade (Supplementary Table 8), where relevant genetic impairments have been documented in schizophrenia (34).

### Effects of betaine supplementation on MAP-induced behavioral sensitization and gene expression

We next examined whether betaine could perform psychotropic action in an animal model of schizophrenia by leveraging behavioral sensitization paradigm, where repeated administration of psychostimulants in rodents can enhance the stimulating effect on locomotor activity. This behavioral paradigm is assumed to model positive symptoms of schizophrenia and the relapse process (35). It is of note that peripheral betaine concentration in schizophrenia was inversely correlated with positive symptom scores of the PANSS (Positive and Negative Syndrome Scale) test (3). Water or betaine (2.17%) was administered 3 weeks before methamphetamine (MAP) injection throughout to the day of MAP challenge (Fig. 5A). In the B6N mice, the challenge injection of MAP elicited enhanced locomotor activities compared to those of the first MAP administration in both water and betaine groups (Fig. 5B). Remarkably, betaine suppressed the degree of elevation in the locomotor activities, which was counted as the differences between the challenge MAP-induced activities and the first MAP-induced activities (*p* < 0.01) (Fig. 5C). In the C3HN mice, repeated MAP injections did not evoke a marked locomotor sensitization (Fig. 5D). Accordingly, the effect of betaine on the suppression of behavioral sensitization was not clear in the C3HN mice (Fig. 5E), demonstrating the different pharmacogenetic profiles between the two strains.

**Figure 5.**
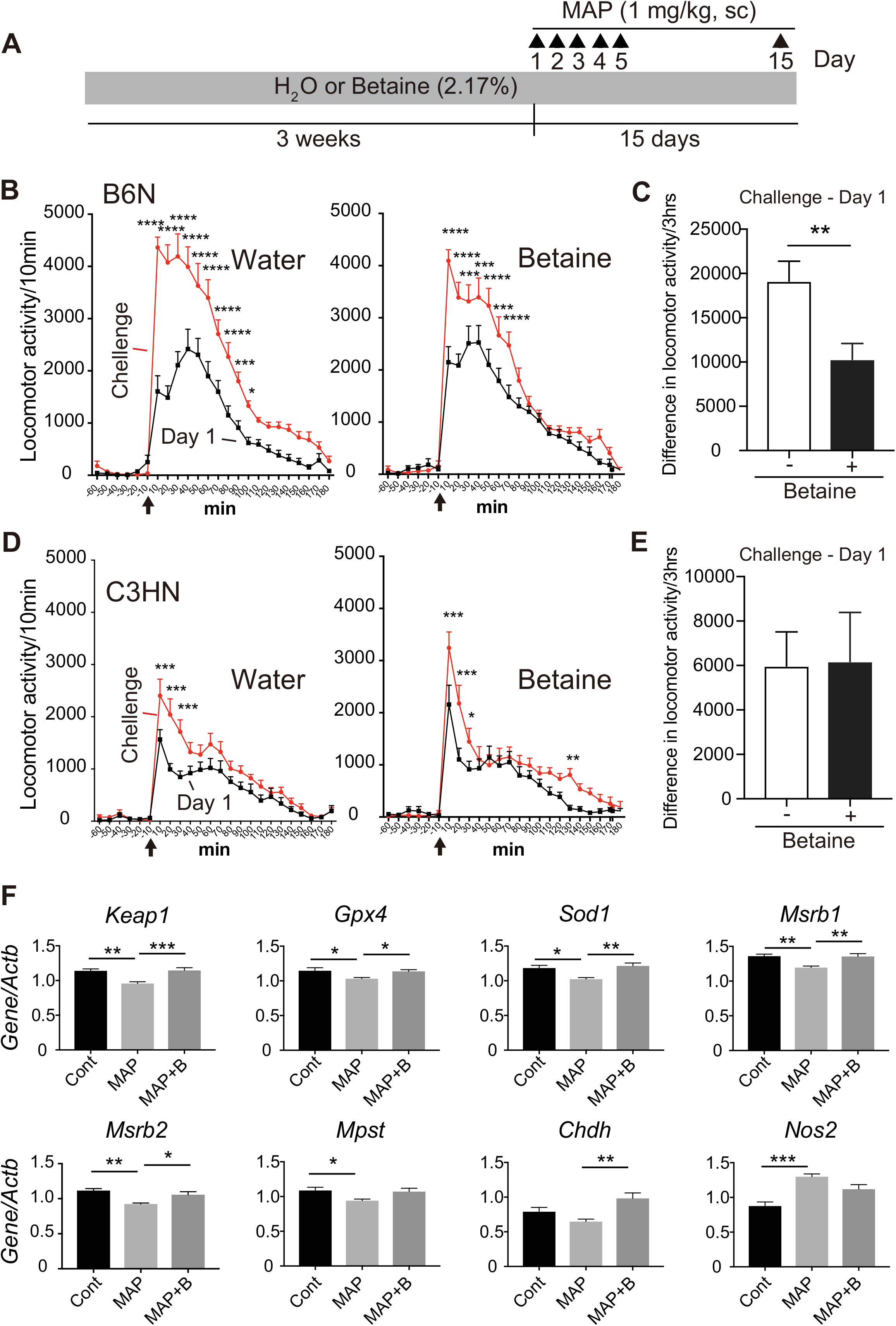
Betaine suppressed methamphetamine-induced behavioral sensitization and restored altered oxidative stress/neuroinflammatory conditions. **(A)** Methamphetamine (MAP) sensitization tests were conducted with (+) or without (−) chronic betaine supplementation in two inbred strains, B6N and C3HN. The activities by first-day injection and challenge injection, and the differences in the locomotor activities “Challenge – Day1” [(locomotor activity induced by challenge injection) – (locomotor activity induced by first-day injection)] are presented **(B–E)**. Betaine supplementation suppressed MAP-induced sensitization in B6N mouse **(B, C)**; whereas, C3HN was not susceptible to the MAP-induced sensitization when compared to B6N **(D, E)**. Data represent mean ± SEM. \**p* < 0.05, \*\**p* < 0.01, *** *p* < 0.001, **** *p* < 0.0001; Bonferroni’s multiple comparison test after repeated ANOVA in **(B, D)**, and Student *t*-test in **(C, E)**; *n* = 12 per group. **(F)** After B6N mice received repeated MAP injections followed by challenge injection, expressions of antioxidant and proinflammatory genes in the frontal cortex were examined. Data represent mean ± SEM. \**p* < 0.05, \*\**p* < 0.01, *** *p* < 0.001; Bonferroni multiple comparison test between two preset pairs: Cont vs. MAP, and MAP vs. MAP+B; *n* = 5–6, per group. Cont, control (H_2_O + repeated saline + MAP challenge); MAP, methamphetamine (H_2_O + repeated MAP + MAP challenge); MAP+B, methamphetamine + betaine (betaine + repeated MAP + MAP challenge).

While the gene expression analyses revealed that betaine deficits and supplementation influenced multiple schizophrenia-relevant transcriptome signatures, we specifically focused on the gene expressions relevant to oxidative stress and neuroinflammation in the MAP model, because (1) such mechanism has been presumed as a pathophysiological component of behavioral sensitization (36) and schizophrenia (37), and (2) betaine’s potency of antioxidant/anti-inflammatory activity is proposed (Supplementary Fig.1) (9). It is suggested that betaine could promote nonenzymatic antioxidant activity through the accelerated turnover of methionine in the methionine–homocysteine cycle. Methionine can act as a scavenger of reactive oxygen species (ROS) (38), and oxidized methionine, methionine sulfoxide, can be reduced back to methionine by methionine sulfoxide reductase A (MSRA), B1 (MSRB1) and B2 (MSRB2) (Fig. 1) (39). Repeated MAP pretreatments coordinately dampened the expressions of antioxidant genes (*Keap1, Gpx1, Gpx4, Sod1, Msrb1, Msrb2* and *Mpst*) (*p* < 0.05) induced by the MAP challenge, and in contrast upregulated the proinflammatory gene *Nos2* (*p* < 0.05), from the comprehensive panel of genes tested (Fig. 5F, Supplementary Table 10 and Supplementary Fig. 8). Betaine cotreatment antagonized the effects of repeated MAP pretreatments on the expression of multiple antioxidant genes (*p* < 0.05 for *Keap1, Gpx4, Sod1, Msrb1* and *Msrb2*; *p* = 0.05 for *Mpst*) (Fig. 5F). Betaine cotreatment also revealed a trend towards lowered response of proinflammatory gene expression (*Nos2, p* = 0.06) after the MAP challenge (Fig. 5F). These results suggest that betaine’s action against MAP-induced sensitization involves, at least in part, antioxidant/proinflammatory response system.

### Altered oxidative stress and proinflammatory conditions were rescued by betaine administration in *in vitro* phencyclidine model

We further pursued betaine’s action against oxidative stress conditions in *in vitro* phencyclidine (PCP) model. Chronic administration of PCP, a noncompetitive *N*-methyl-d-aspartate (NMDA) receptor blocker, produces schizophrenia-like behaviors in humans (40). This psychotomimetic effect is known to be partly associated with oxidative stress given by PCP (41). Because the deteriorating effect of PCP on neuronal system is also observed in the primary neuron culture (42), we examined the expressional changes of the same genes as in the MAP treatment experiments, plus *Sod3* (this gene is expressed in rat brain but not in mouse brain), in this paradigm (Fig. 6A). Among those genes, the expressions of *Nfe2l2, Rela, Cat, Gpx1, Sod2, Sod3, Cth, Tnf, NFkb1* and *Nos2* showed coordinate upregulation (*p* < 0.05) under the PCP (1 μM) exposure (Fig. 6B, also see Supplementary Fig. 9). And notably, the addition of betaine (500 μM) led to significant expressional suppression (Fig. 6B). These results demonstrate again a role of betaine in the antioxidant and proinflammatory system.

**Figure 6.**
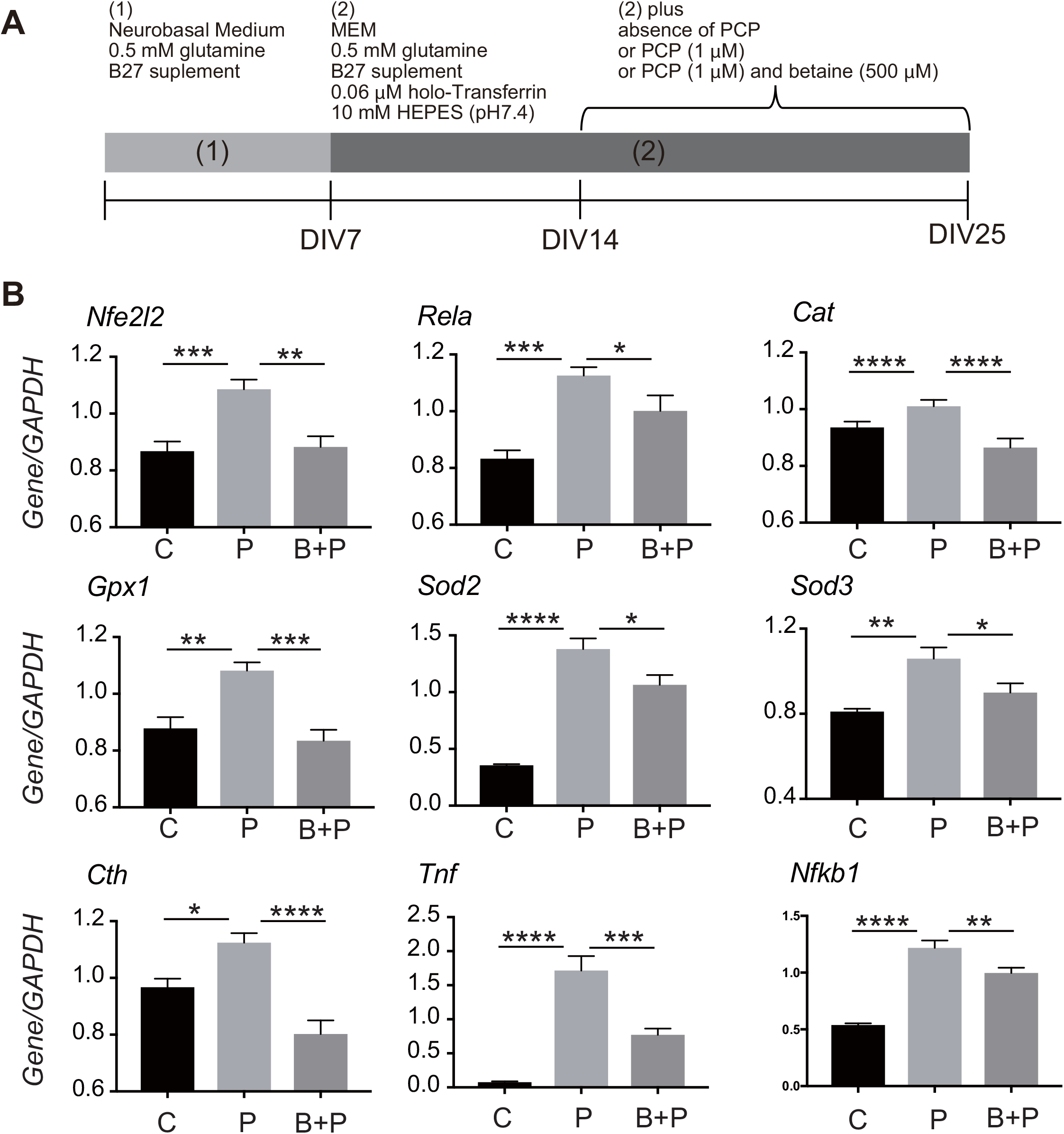
Betaine alleviates PCP-induced pro-inflammatory and antioxidant gene expressional changes in primary cortical neurons. **(A)** After the preparation and incubation of rat cortical neurons, they were maintained in the absence of PCP, or in the presence of PCP (1 μM) or PCP (1 μM) plus betaine (500 μM) for 11 days. **(B)** Total RNAs were prepared for real time RT-PCR to examine expression of each gene (see the text for the details). Data represent mean ± SEM. \**p* < 0.05, \*\**p* < 0.01, \*\*\**p* < 0.001, \*\*\*\**p* < 0.0001; Tukey’s multiple comparison test. C; control (absence of PCP), P; PCP, B+P; betaine + PCP.

### Analyses of betaine levels and carbonyl stress in postmortem brain samples

Next, we asked whether betaine-related pathology can be seen in schizophrenia. Interestingly, reduced levels of betaine were observed in the postmortem brain tissue from patients with schizophrenia when compared to the controls (Fig. 7A), while the other metabolites unchanged (Figs. 7B-G) (16–18) (Supplementary Table 11). SAM contents showed a decreased trend (*p* < 0.1), whereas SAH levels were unaltered in the schizophrenia group, resulting in reduced SAM/SAH ratio in the schizophrenic brain (*p* < 0.01) (Figs. 7H-J), which can be linked to the decreased betaine levels (Fig. 1). The betaine levels were not affected by the confounding factors (Supplementary Table 12).

**Figure 7.**
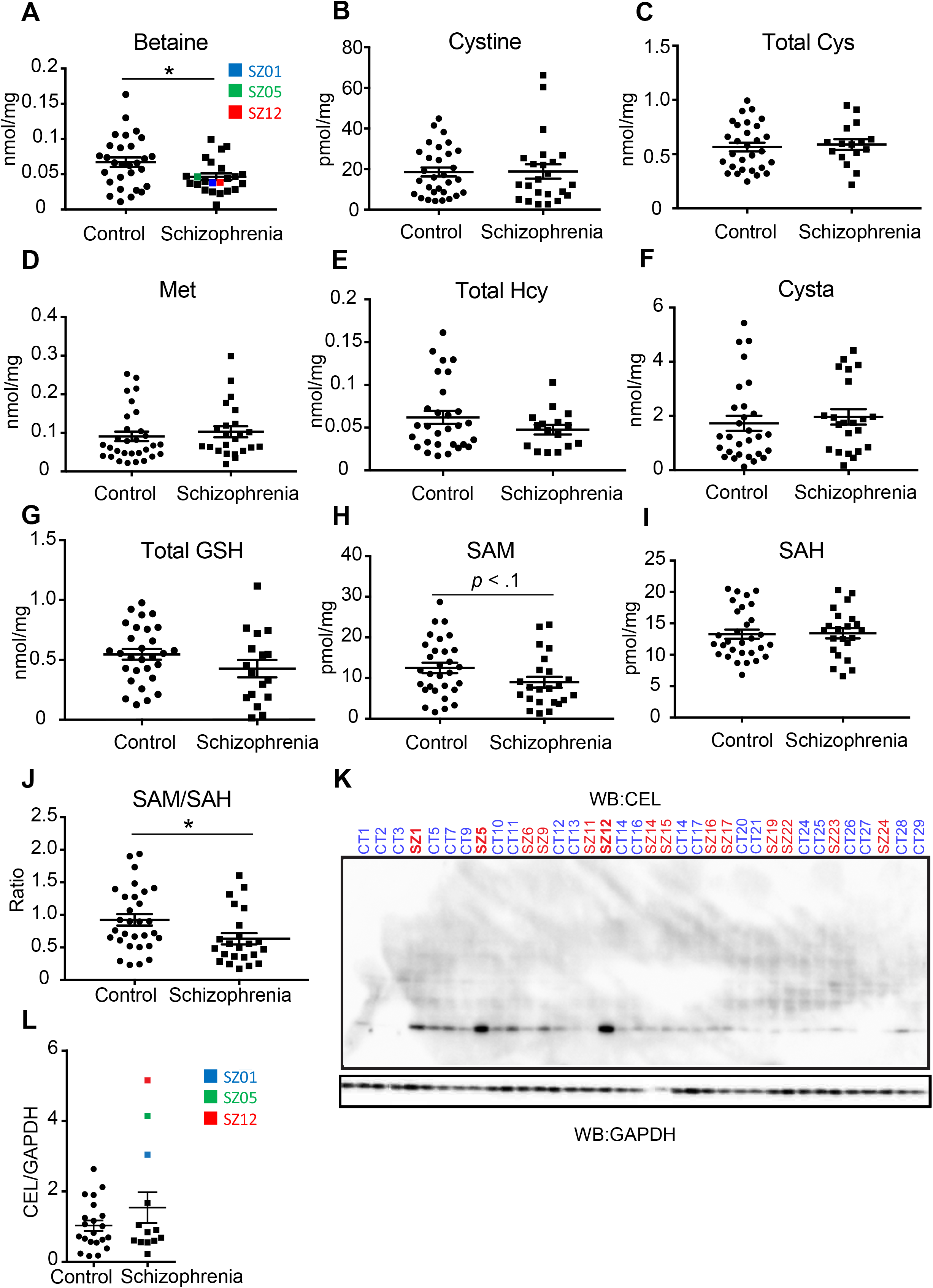
Measurements of metabolite contents and advanced glycation end products (AGEs) in postmortem brain samples from schizophrenia. **(A)** Betaine levels are significantly reduced in postmortem schizophrenic brain samples. No other metabolites such as **(B)** cystine, **(C)** total Cys, **(D)** methionine (Met), **(E)** total Hcy, **(F)** cystathionine (Cysta), **(G)** total glutathione (GSH + GSSG), **(H)** *S*-adenosylmethionine (SAM), and **(I)** *S*-adenosylhomocysteine (SAH) were significantly different between schizophrenia cases and controls. **(J)** The ratio of SAM to SAH (SAM/SAH) was decreased in schizophrenia aligning with reduced betaine levels. Data represent mean ± SEM. \**p* < 0.05; two-tailed Student’s *t*-test. *n* = 29–31 in the control group, *n* = 22–23 in the schizophrenia group, except for total GSH, total Cys, and total Hcy measurements, where *n* = 16–17 in the schizophrenia group was used. **(K, L)** Three postmortem brain samples (SZ1, SZ5, and SZ12) from patients with schizophrenia revealed enhanced signals by anti-CEL antibody, while no statistical difference was seen between control and schizophrenia groups **(L)**. Data represent mean ± SEM. \**p* < 0.05; two-tailed Student’s *t*-test. *n* = 22 in the control group, *n* = 14 in the schizophrenia group.

To investigate the relationship between betaine system abnormality and oxidative stress conditions in schizophrenia, we specifically focused on the carbonyl stress in the postmortem brain samples. AGEs analysis identified the three schizophrenia brain samples (#SZ01, #SZ05 and #SZ12) that showed a strong band (~30 kDa protein(s)) having Nε-(carboxyethyl)lysine (CEL) (Figs. 7K and L), one of AGEs (13). The three samples showed betaine levels of below average (Fig. 7A), suggesting a relationship between elevated carbonyl stress and lowered betaine levels. Furthermore, these three patients displayed relatively high symptom scores evaluated by using the Diagnostic Instrument for Brain Studies (DIBS) (43) (Supplementary Table 13). Collectively, the results of postmortem brain study highlighted the existence of a subset of patients with schizophrenia characterized by “betaine deficit-oxidative stress” pathology, manifesting relatively severe psychotic symptoms.

### Betaine’s potency in alleviating carbonyl stress in *GLO1*-deficient hiPSCs

Because coexistent betaine pathology and elevated carbonyl stress were evident in a subset of schizophrenia cases, the efficacy of betaine supplementation in mitigating the carbonyl stress in isogenic lines of *GLO1*-deficient hiPSCs was tested. We firstly confirmed that there were no missense variants in the genome of the subject (mentally healthy) for the genes in the carbonyl stress pathway (Supplementary Fig. 2). The *GLO1*-deficient hiPSC lines were prepared by harnessing the CRISPR-Cas9 system, and they contained homozygous *GLO1* KO alleles (KO-1, KO-2 and KO-3) causing frameshift mutations in exon 1 and premature stop signals (Figs. 8A and B), and showed no protein expression (Fig. 8C). As controls, three lines (WT-1, WT-2 and WT-3) were established that were proven to harbor the normal *GLO1* alleles. The *GLO1*-deficient hiPSC samples displayed a major band of ~55 kDa (called “band A”) with increased modification by carboxymethyllysine (CML), another species of AGEs (13) (representative picture in Fig. 8C). When betaine was added to cell culture, concentration at 500 μM significantly suppressed the CML modification of “band A” (Fig. 8D), indicating betaine’s efficacy against elevated carbonyl stress. The *GLO1*-disrupted hiPSCs showed comparable morphology (Fig. 8B) with those of the WT, but they manifested reduced differentiation potential into neural lineage, which could not be rescued by the betaine supplementation (data not shown).

**Figure 8.**
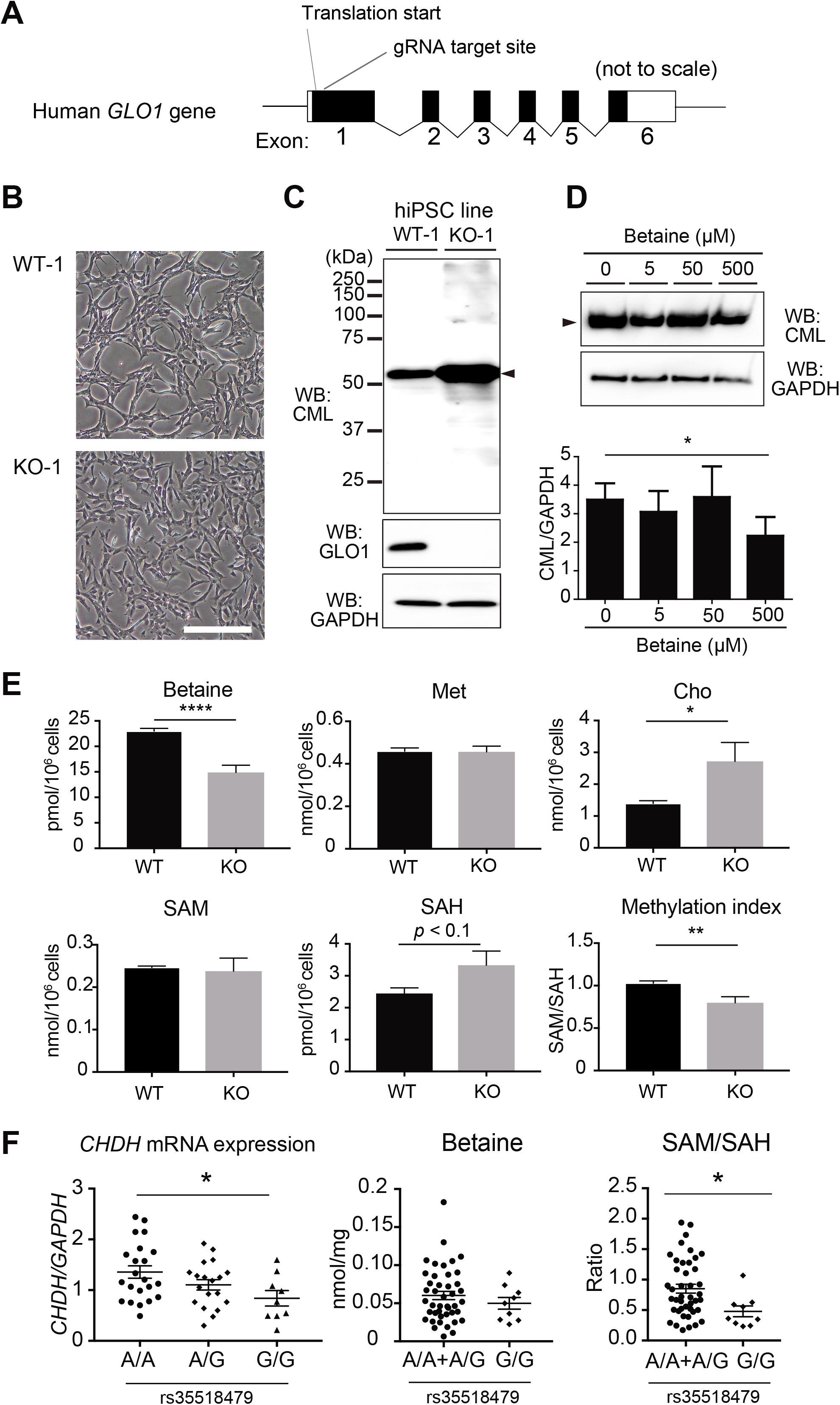
Betaine suppressed accumulation of advanced glycation products (AGEs) in hiPSCs and a genetic variant affected *CHDH* expression in brain. **(A)** Genomic structure of the human *GLO1* gene is schematically presented. Black and white boxes indicate coding and untranslated regions of exons, respectively. Three hiPSC lines for each genotype (WT-1-3 and KO-1-3) were established. **(B)** Phase contrast images of WT-1 and KO-1 are presented. WT-2-3 and KO-2-3 showed morphology similar to WT-1 and KO-1 (data not shown). Scale bar: 400 μm. **(C)** Cell (WT-1 and KO-1) lysates were analyzed in western blotting using an antibody against carboxymethyllysine (CML), a typical type of AGEs. Note that the signal around 55 kDa (arrow head; “band A”) was greatly enhanced by depletion of the intact GLO1 proteins. The filter was also probed with anti-GAPDH antibody as a loading control and anti-GLO1 antibody to show a loss of the GLO1 protein in the *GLO1* KO (−/−) cells. **(D)** KO-1 was cultured for four days in the absence or presence of indicated concentrations (5, 50, or 500 μM) of betaine. The results of western blot analysis of the cell lysates are presented (top). The position for “band A” is indicated by an arrowhead. The same experiment was repeated using KO-2-3 and WT-1-3 in duplicate. A significant difference (*n* = six per group; 3 KO or WT lines × duplicate) was observed between the groups treated with 0 μM and 500 μM betaine. The CML signals were normalized to the intensity of GAPDH signals. Data represent mean ± SEM. \**p* < 0.05; Dunnett multiple comparison test. *n* = six. **(E)** Contents of metabolites in the methionine-homocysteine cycle in the *GLO1* KO hiPSCs. Data represent mean ± SEM. \*\**p* < 0.01, \*\*\**p* < 0.001; two-tailed Student’s *t*-test. WT-1-3 and KO-1-3 were triplicated (*n* = nine each). **(F)** *CHDH* variant, rs35518479, showed a significant association for its expression and SAM/SAH ratio in the brain, though betaine levels were unaltered. Data represent mean ± SEM. \**p* < 0.05, \*\**p* < 0.01, \*\*\**p* < 0.001; one-way ANOVA and two-tailed Student’s *t*-test.

Next, we examined the effects of *GLO1* ablation on the levels of betaine and metabolites in the methionine-homocysteine cycle. Strikingly, betaine levels were significantly decreased in the *GLO1* KO hiPSCs compared to the WT cells (Fig. 8E), suggesting that betaine is consumed to cope with increased carbonyl stress. In parallel, lowered SAM/SAH ratio concomitant with increased SAH levels in the KO cells was seen (Fig. 8E), probably reflecting dampened intracellular homocysteine clearance by BHMT pathway (Fig. 1) in the KO cells.

### Pharmacogenetic evaluation of betaine efficacy in schizophrenia

We have revealed the differential efficacy for betaine in the mouse strains with distinct genetic background (Fig. 5). Since a subset of schizophrenia patients were characterized with betaine deficit-oxidative stress pathology and the betaine treatment was effective in alleviating carbonyl stress, we speculated that the efficacy for betaine might have a genetic predisposition. *Cis*-eQTL analysis of common genetic variants in *BHMT1, CHDH* and *GLO1* genes in postmortem brain tissues revealed a significant association of a *CHDH* variant, rs35518479, with its expression levels in the brain (*p* = 0.03) (Fig. 8F, Supplementary Figs. 10A, B). The A allele carriers showed a significantly higher *CHDH* expression levels in the brain when compared to the G/G homozygotes. Although there were no significant differences in betaine and SAM levels according to the genotypes (Supplementary Fig. 10 C), the SAM/SAH ratio (methylation index) was higher in the A allele carriers (*p* = 0.02) (Fig. 8F). The results suggest that *CHDH* expression levels may affect the turnover rate of methionine-homocysteine cycle, and depict a potential utility of *CHDH*-eQTL in stratifying schizophrenia patients for betaine treatment.

## Discussion

In the present study, we provided a proof of concept for the potential of betaine in the treatment of schizophrenia from the analyses of both mouse and human samples, along with the identification of “betaine deficit-oxidative stress” pathology in a subset of schizophrenia, which may result in relatively sever symptoms. Our systematic examination of betaine’s role (see Supplementary Fig. 1) showed that the betaine’s action as a psychotropic is imparted partially through antioxidant/proinflammatory effects. In addition, a regulation of DNA methylation capacity may be possible, from metabolomics/methylome analyses of mouse and human samples. Notably, analyses of two different inbred mouse strains highlighted an interaction of genetic background × drug (e.g. betaine), and distinct pharmacogenetic profiles (e.g. MAP) according to genetic background.

In this study, we detected a substantial accumulation in CEL adduct of ~30 kDa protein(s) (Fig. 7K) and CML adduct of ~55 kDa protein(s) (Fig. 8C) in postmortem brain and *GLO1*-KO iPSCs, respectively. It was unexpected to see AGEs in a relatively protein-specific manner under carbonyl stress, since the Maillard reaction (Supplementary Fig. 2) is deemed to occur non-enzymatically (13). Molecular identification and characterization of the specifically modified proteins and the mechanism of the specific modifications remain to be studied.

Although we did not address in this study, another known function of betaine is osmotic regulation (Supplementary Fig. 1), and the deficits in brain betaine levels may contribute to cellular osmotic perturbation (7, 8, 44), which is reported to inhibit methionine uptake, inhibit protein synthesis, and affect the mRNA translation by dysregulation of phosphorylation and mTOR signaling cascades (45–47). These pathways are shared with those identified in the transcriptomics analysis of *Chdh*-deficient mice (Fig. 3F). Although taurine, *myo*-inositol, glycine and glutamine are thought to play major roles as osmolytes in the brain because of their higher cellular contents, betaine is favored in neural cells as an osmolyte (7). Thus, the osmotic issue should be pursued in future studies.

Since betaine has already been approved as a therapeutic drug for homocystinuria, an autosomal recessive inherited disorder due to a deficiency of cystathionine beta synthase (CBS, Fig. 1) (48), it has the advantage of being repositioned for psychiatric illness. The utility is also substantiated by our observation that betaine can penetrate the blood-brain barrier and is well tolerated with no serious adverse effects.

Interestingly, we identified an eQTL for *CHDH* expression, which substantiates genotype-based stratification of schizophrenia patients for betaine efficacy. Future studies are warranted to validate the utility of “betaine deficit-oxidative stress pathology” as a biomarker and to identify other genetic underpinnings for betaine’s efficacy. It is also worthwhile to test the availability of betaine in clinical setting for schizophrenia and other neuropsychiatric conditions.

## Acknowledgements

We are grateful to the Support Units for Bio-Material Analysis and Animal Resources Development at RIKEN CBS Research Resources Division, particularly Mr. Masaya Usui and Mr. Hiromasa Morishita for the development of the LC/MS and HPLC systems to measure the contents of various metabolites, and Mr. Takashi Arai for the technical supports in generating the *Chdh*-deficient mice. We also appreciate Ms. Chiaki Watanabe and Hiromi Onuma (Postmortem Brain Bank of Fukushima for Psychiatric Research) for their contribution in sample collection coordination. Eventually, we wish to express special thanks to the families of the deceased for the donations of brain tissue, and for their time and effort devoted to the consent process and interviews.

## Supplementary materials

Supplementary materials are available online.

