## Supplementary Fig. for "Investigation of betaine as a novel psychotherapeutic for schizophrenia"

### Supplementary Figures

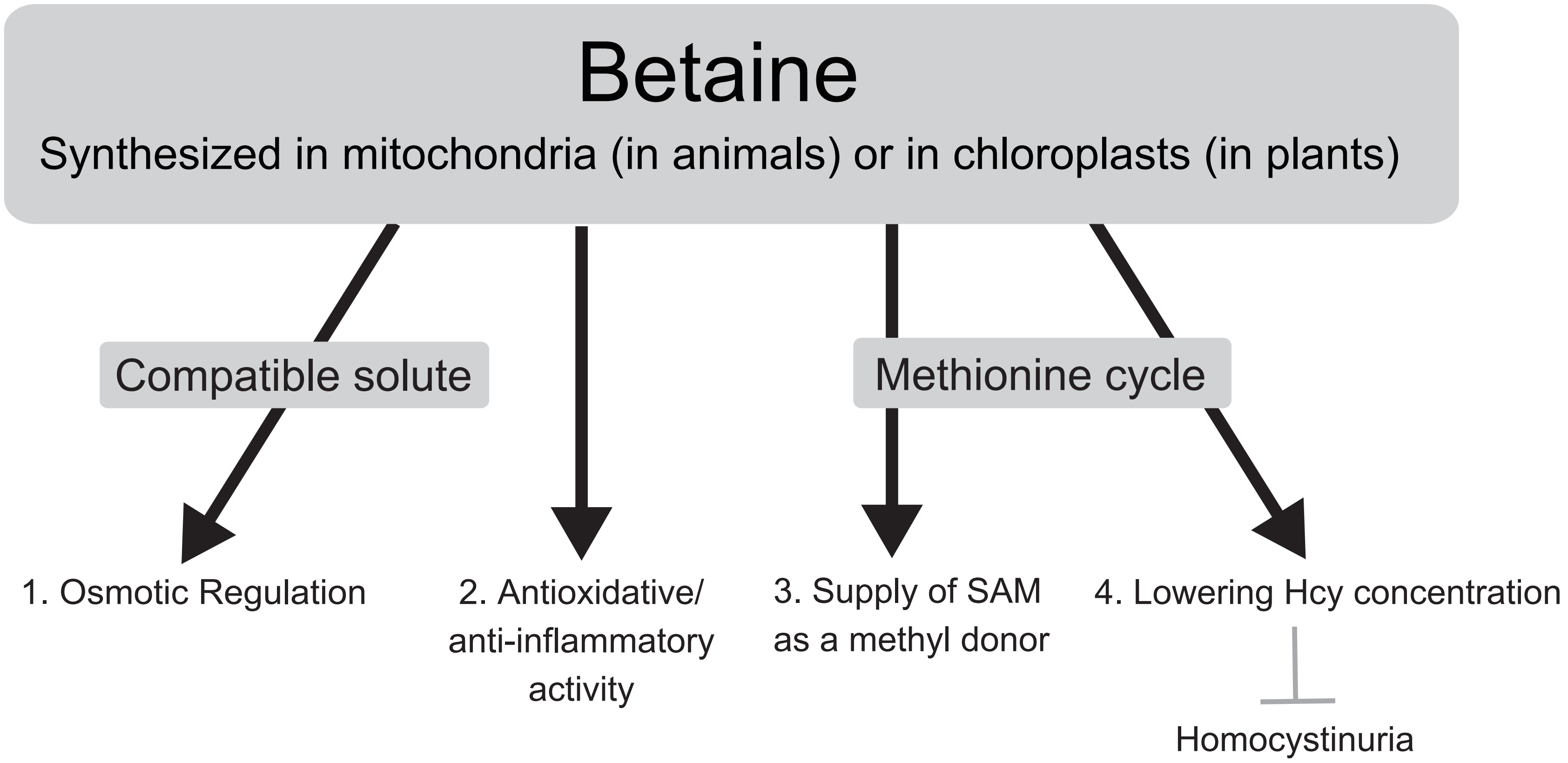

**Supplementary Figure 1. Biological functions of betaine**. Betaine is thought to act as osmotic regulator (compatible solute), provider of antioxidative/anti-inflammatory activity, supplier of methyl donor S-adenosylmethionine,and mitigator of elevated homocysteine levels.

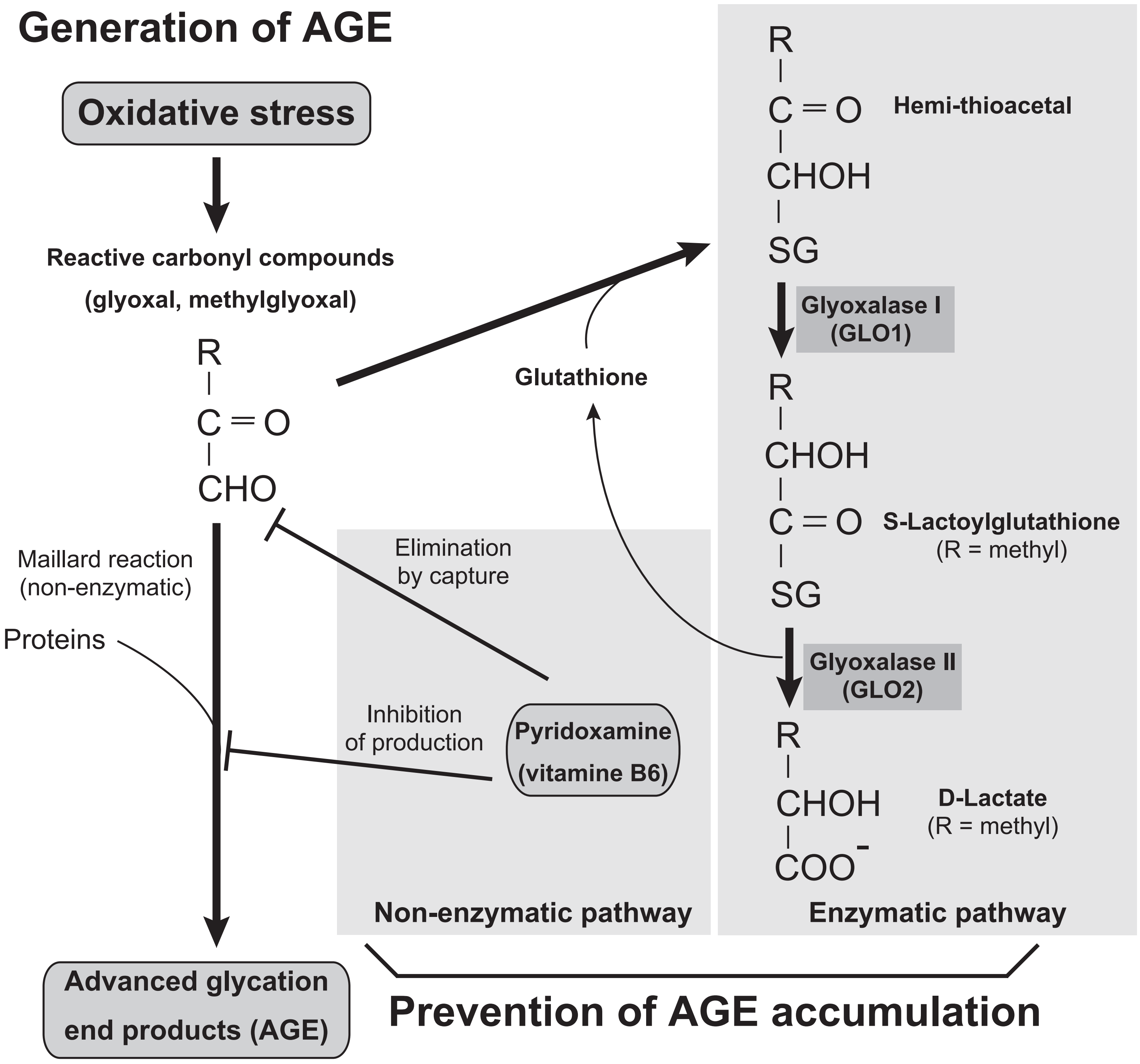

**Supplementary Figure 2. Biochemical pathways involved in the production of advanced glycation end products (AGEs).** The zinc metalloenzyme glyoxalase 1(GLO1) and nonenzymatic action of pyridoxamine are critically involved in the prevention of cellular accumulation of AGEs. The buildup of AGEs is known to underlie the pathophysiology in a subset of schizophrenia.

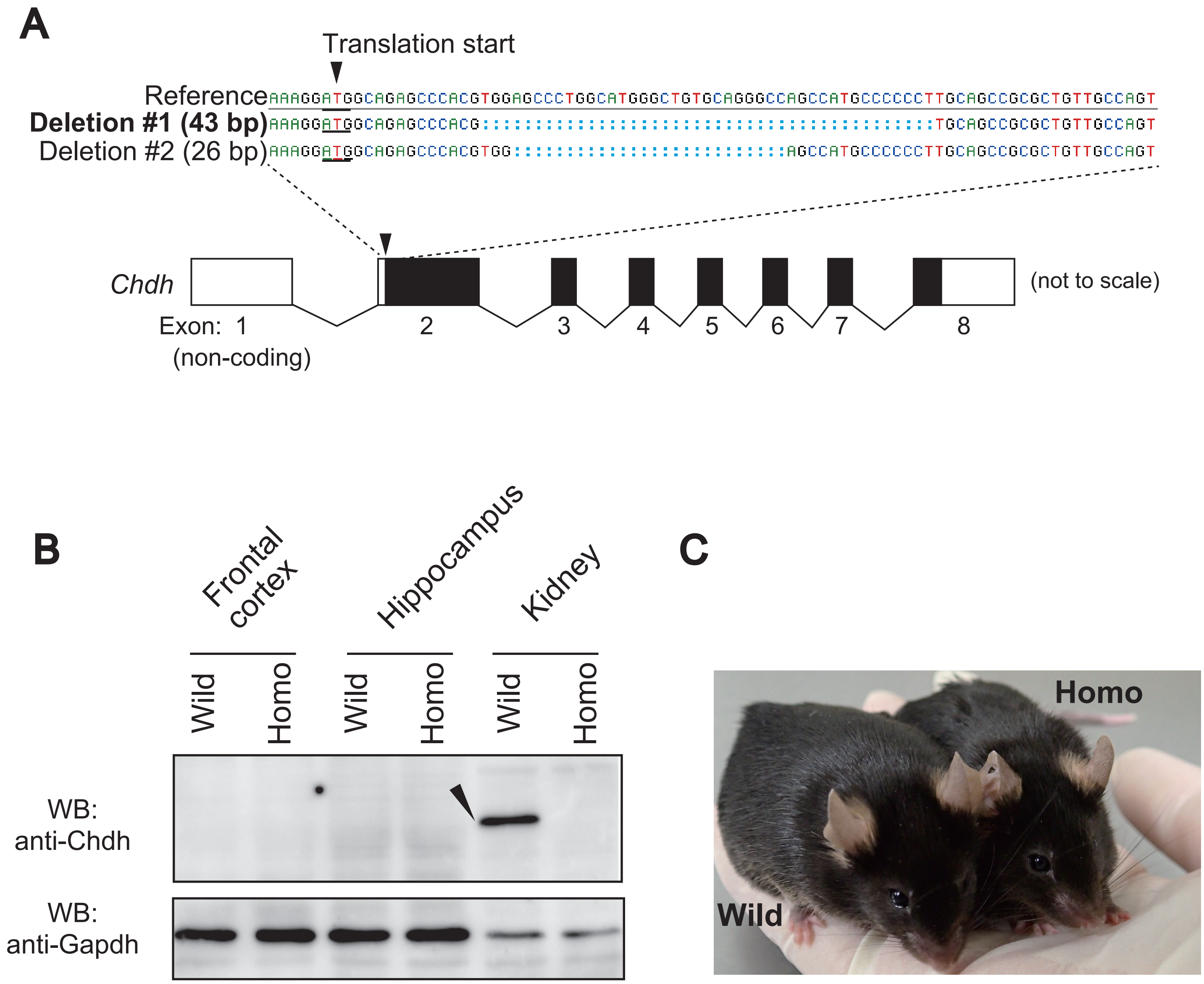

**Supplementary Figure 3. Generation of *Chdh*-null mice.** (**A**) Gene editing of *Chdh* by the CRISPR/Cas9n methodology. Schematic of the genomic structure of the mouse *Chdh* gene is presented. Black and white boxes and exons of the gene indicate coding and untranslated regions, respectively. The founder #509 harbored two different deletions (deletion#1 and deletion #2) in the first coding exon (exon 2) of *Chdh*. (**B**) Western blot analysis using anti-Chdh antibody. A discrete band (arrow head) was detected in the kidney from wild-type, but not in homozygote for deletion #1. Note that two brain regions (frontal cortex and hippocampus) do not reveal expression levels sufficient to detect specific signals. (**C**) No gross abnormalities were seen in homozygotes.

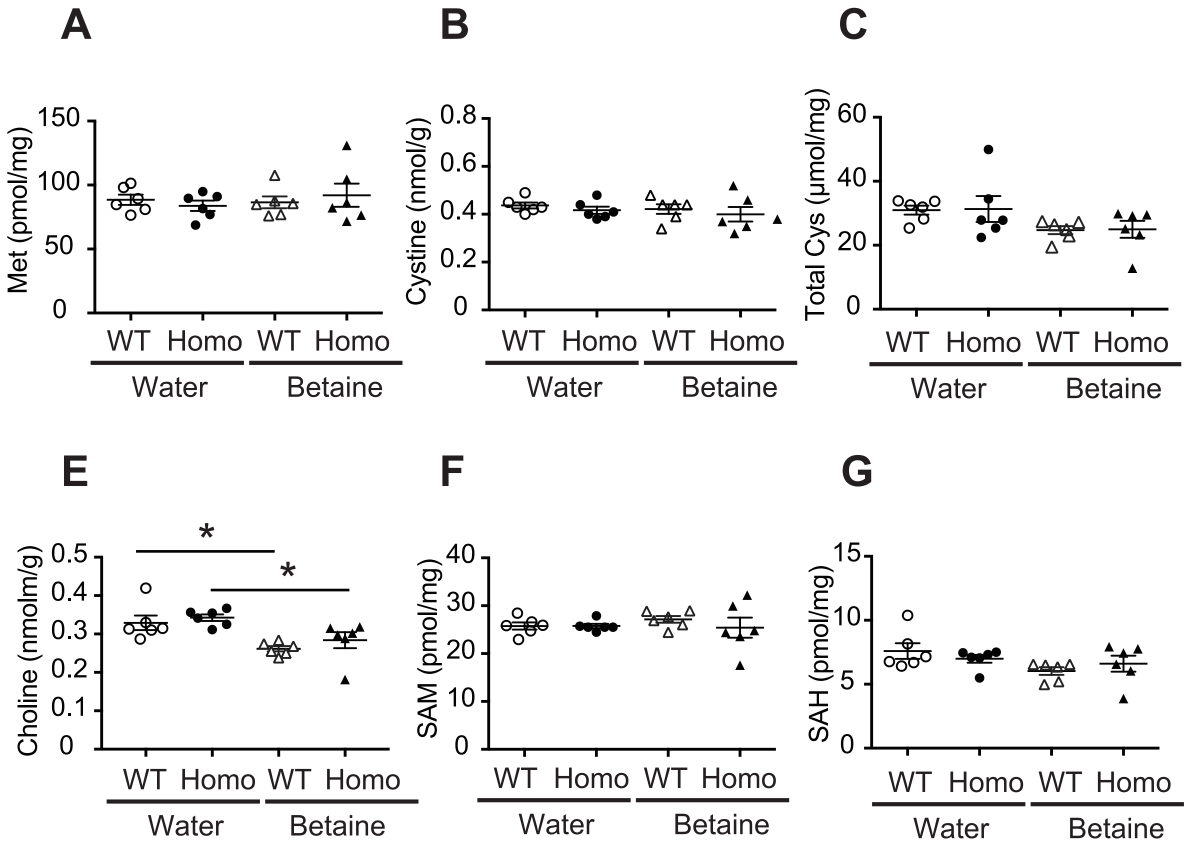

**Supplementary Figure 4. Effect of a functional loss of *Chdh* gene and betaine supplementation on metabolite contents in the frontal cortex**. The metabolites (**A**) methionine, (**B**) cystine, (**C**) total cysteine, (**D**) cystathionine, (**E**) choline, (**F**) *S*-adenosylmethionine (SAM), and (**G**) *S*-adenosylhomocysteine (SAH) in the brain (frontal cortex) were not affected by the gene disruption or betaine supplementation. Data represent mean ± SEM. *$P < 0.05$, **$P < 0.01$; unpaired two-tailed Student' s *t*-test. *n* = 6–7 per group. Homo, homozygous *Chdh*-deficient mouse; WT, wild-type littermate controls.

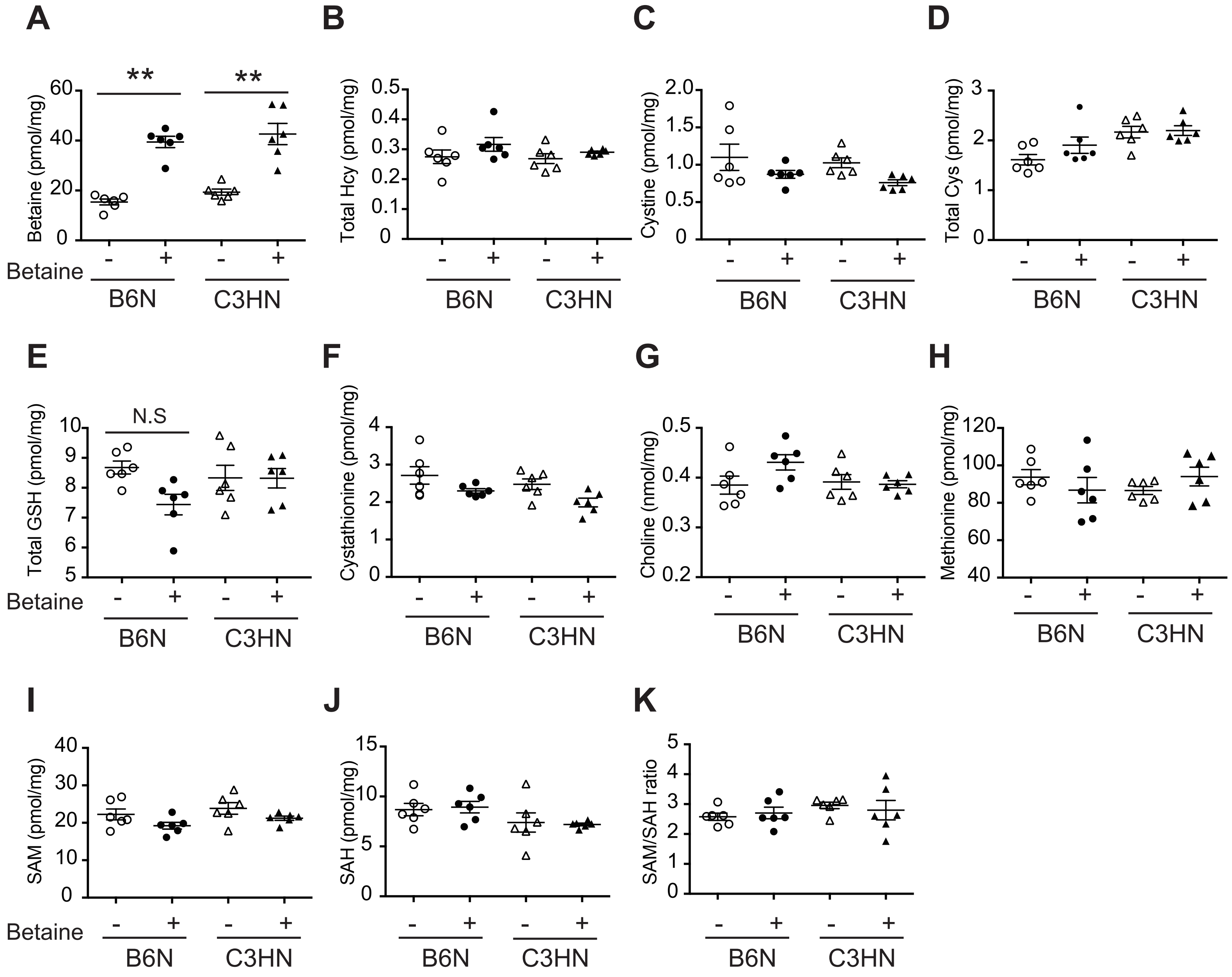

**Supplementary Figure 5. Effect of betaine supplementation on metabolite contents in the frontal cortex from two inbred mouse strains, B6N and C3HN.** Concentration of metabolites (**A**) betaine, (**B**) total Hcy, (**C**) cystine, (**D**) total Cys, (**E**) total GSH (GSH + GSSG), (**F**) cystathionine, (**G**) choline, (**H**) methionine, (**I**) *S*-adenosylmethionine (SAM), (**J**) S-adenosylhomocysteine (SAH), and (**K**) SAM/SAH ratio in the frontal cortex of inbred mouse strains upon betaine supplementation. Betaine levels were increased in frontal cortex upon supplementation in both inbred strains of mice, indicating that betaine can cross the blood-brain barrier. Data represent mean ± SEM. *$p < .05$, **$p < .01$; unpaired two-tailed Student' s *t*-test. $n$ = 5–6 per group.

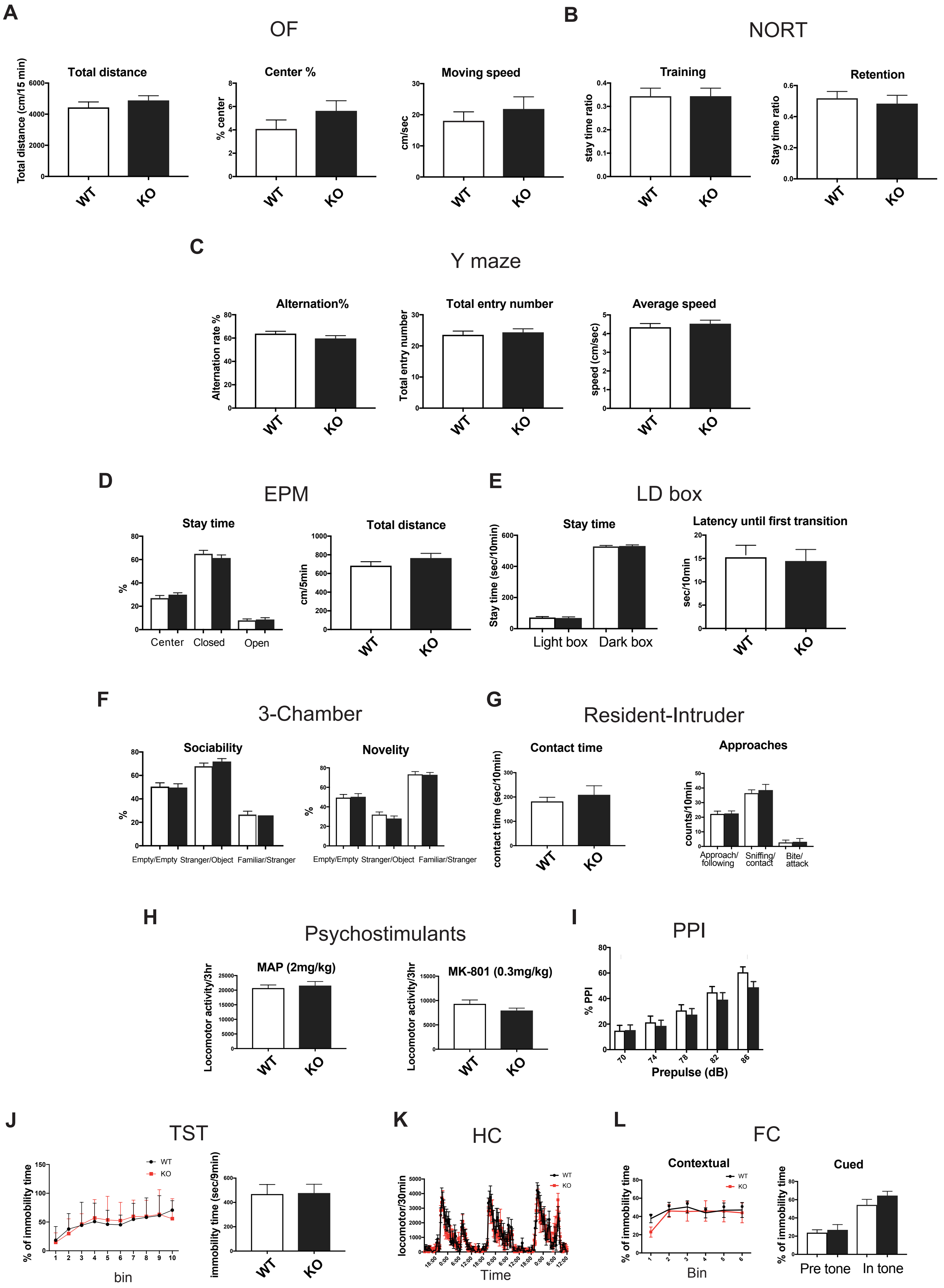
A
OF
Total distance
Total distance (cm/15 min)
Center %
% center
Moving speed
cm/sec
B
NORT
Training
stay time ratio
Retention
Stay time ratio
WT
KO
C
Y maze
Alternation%
Alternation rate %
Total entry number
Total entry number
Average speed
speed (cm/sec)
D
EPM
Stay time
%
Center
Closed
Open
Total distance
cm/5min
E
LD box
Stay time
Stay time (sec/10min)
Light box
Dark box
Latency until first transition
sec/10min
F
3-Chamber
Sociability
Novelity
%
Empty/Empty
Stranger/Object
Familiar/Stranger
G
Resident-Intruder
Contact time
contact time (sec/10min)
Approaches
counts/10min
Approach/ following
Sniffing/ contact
Bite/ attack
H
Psychostimulants
MAP (2mg/kg)
MK-801 (0.3mg/kg)
Locomotor activity/3hr
I
PPI
% PPI
Prepulse (dB)
J
TST
% of immobility time
bin
immobility time (sec/9min)
K
HC
locomotor/30min
Time
L
FC
Contextual
Cued
% of immobility time
Bin
Pre tone
In tone

**Supplementary Figure 6. Behavioral analyses of *Chdh*-deficient mice.** Homozygotes (KO) were tested for a battery of behavioral tests relevant for psychiatric disorders in comparison to wild-type (WT) littermate controls (**A**–**L**). None of the test revealed significant differences between the two genotypes. (**A**) Open field test (OF), $n = 20$ per group, (**B**) Novel object recognition test (NORT), $n = 20$ per group, (**C**) Y-maze test, $n = 20$ per group, (**D**) Elevated plus maze (EPM) , $n = 20$ per group, (**E**) Light-dark box transition test (LD box) , $n = 10$ per group, (**F**) Three-chamber test (3-Chamber), $n = 16$ per group, (**G**) Resident intruder test, $n = 4$ and 5 for WT and homo, respectively, (**H**) Locomotor activities after MAP (ip; 2 mg/kg, left) or MK-801 (ip; 0.3 mg/kg) injection, $n = 8$ per group, (**I)** Prepulse inhibition (PPI) test, $n = 20$ per group, (**J**) Tail suspension test (TST), $n = 10$ per group, (**K**) Home cage activity for 3 days (HC), $n = 8$ per group, and (**L**) Fear conditioning test, contextual (left) and cued (right) tests, $n = 8$ per group. Data represent mean ± SEM. *$P < 0.05$, **$P < 0.01$; unpaired two-tailed Student's *t*-test.

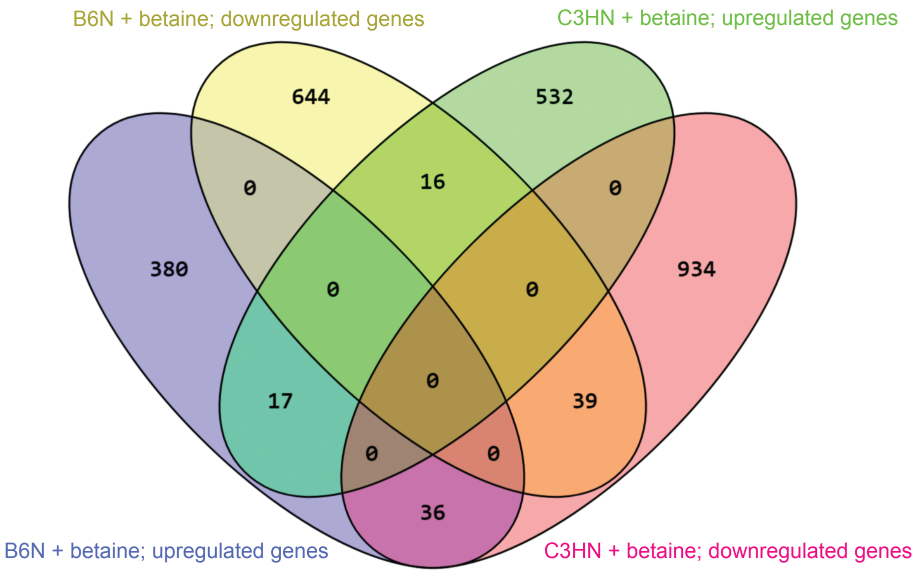

**Supplementary Figure 7.** The venn diagram shows the overlap of transcripts among the differentially expressed genes between B6N and C3HN mice administered with betaine.

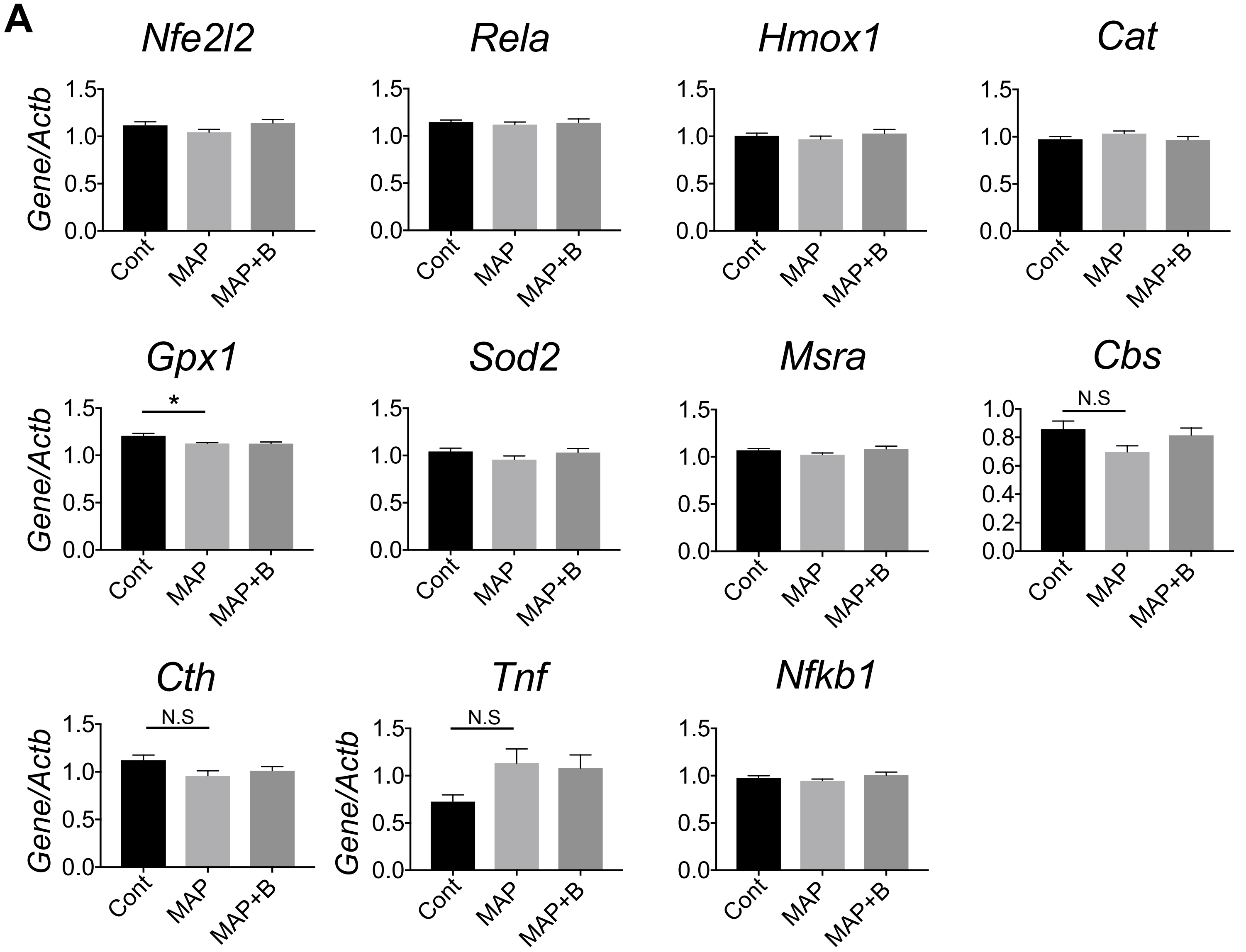

**Supplementary Figure 8. Effects of MAP administration on antioxidant and antiinflammatory genes.** After B6N mice received repeated MAP injections followed by challenge injection, (**A**) expressions of antioxidant and proinflammatory genes in the frontal cortex were examined. Data represent mean ± SEM. *$P < 0.05$; Bonferroni multiple comparison test between two preset pairs: Cont vs. MAP, and MAP vs. MAP + B. $n = 5–6$, per group. Cont, control; MAP, methamphetamine; MAP+B, methamphetamine + betaine.

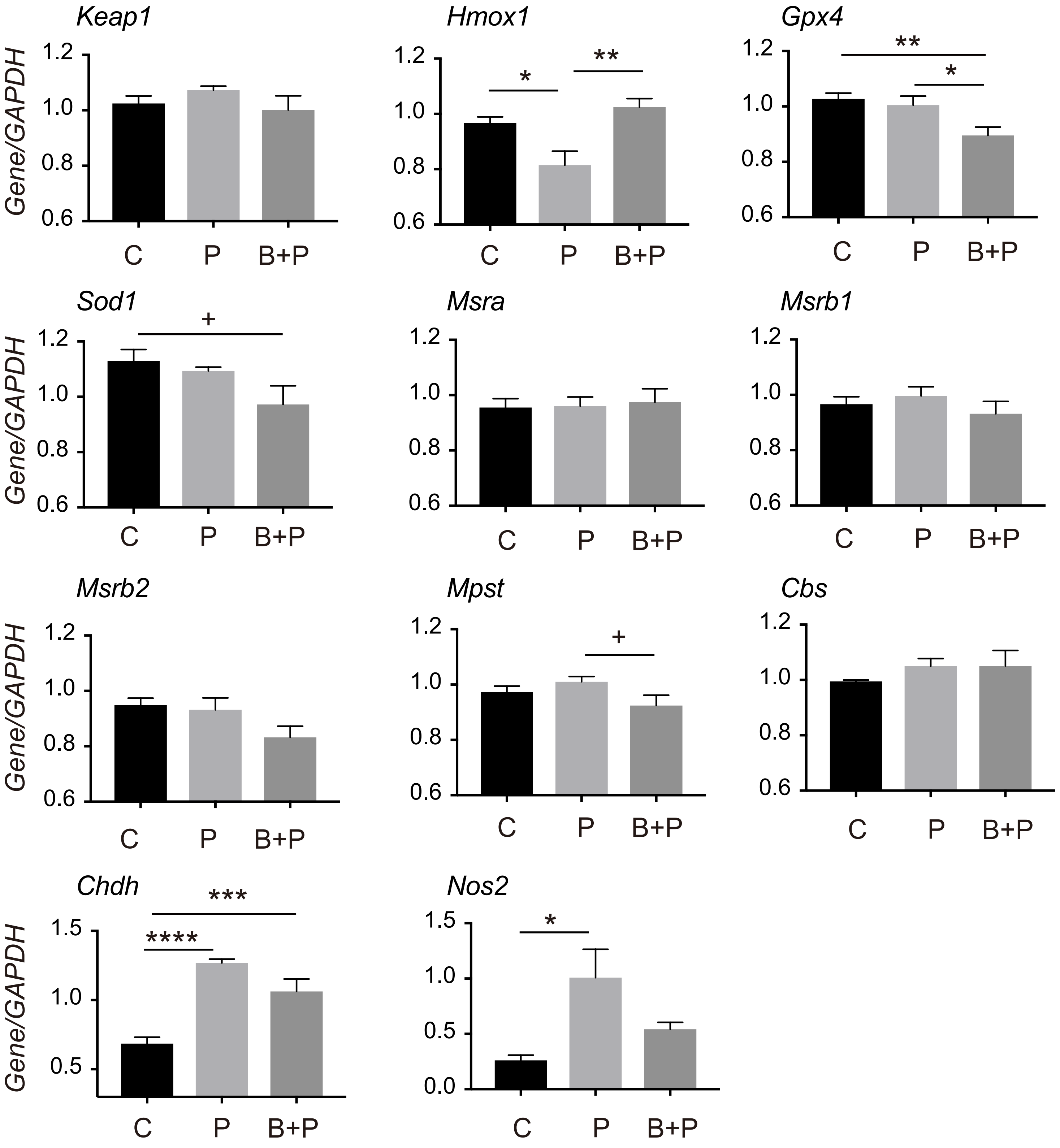

**Supplementary Figure 9. Role of betaine in PCP-induced pro-inflammatory and antioxidant gene expressional changes in primary cortical neurons.** Rat cortical neurons were maintained in the absence of PCP, or in the presence of PCP (1 μM) or PCP (1 μM) plus betaine (500 μM) for 11 days. Total RNAs were prepared for real time RT-PCR to examine expression of each gene (see the text for the details). Data represent mean ± SEM. *$p < 0.05$, **$p < 0.01$, ***$p < 0.001$, ****$p < 0.0001$; Tukey's multiple comparison test. C; control (absence of PCP), P; PCP, B+P; betaine + PCP.

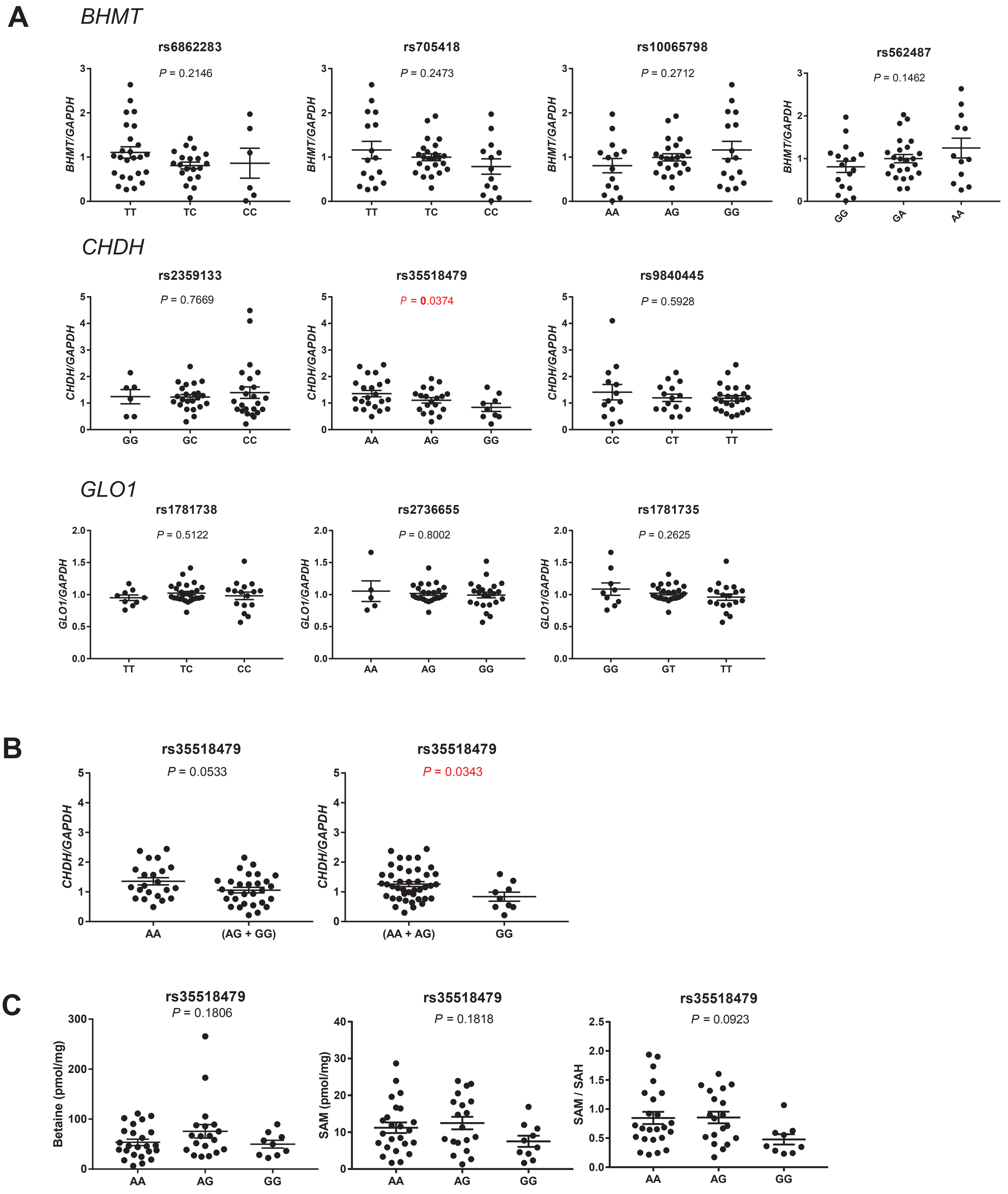

**Supplementary Figure 10. Association analysis of common genetic variants in *BHMT1, CHDH* and *GLO1* with gene expression (cis eQTL) and metabolite levels in postmortem brain tissues**. (**A**) *CHDH* variant, rs35518479, showed a significant association for its expression in the brain and (**B**) the G allele carriers showed significantly low *CHDH* expression levels in the brain when compared to the A allele. (**C**) Levels of betaine, SAM, and SAM/SAH ratio in rs35518479 genotype carriers. Data represent mean ± SEM. *$p < 0.05$, **$p < 0.01$, ***$p < 0.001$; one-way ANOVA and two-tailed Student's t-test. $n = 53$.

### Supplementary methods

#### Study approval

All the animal experiments were performed in compliance with relevant laws, and guidelines were approved by the Animal Ethics Committee at RIKEN Center for Brain Science, Japan (H29-2-204(3), 2016-058(4)). Human induced pluripotent stem cell study was approved by the Human Ethics Committee at RIKEN, Japan for iPSC study (Wako-daisan 25-14). Experiments in postmortem brain samples were approved by Fukushima Medical University, Japan (1685 and 2381) and Niigata University School of Medicine, Japan (G2015-0827).

#### Generation of Chdh-deficient mice by genome editing

The *Chdh*-deficient mice were generated by the genome editing methodology using the CRISPR/Cas9 nickase [1, 2]. Briefly, B6N zygotes obtained by *in vitro* fertilization were microinjected with the cocktail [5 ng/mL Cas9 nickase mRNA (System Biosciences, Mountain View, CA) and 5 ng/mL each two sgRNAs]. The sgRNAs [*Chdh*-upstream (target sequence: 5′- GCCCTGCACAGCCCATGCCAGGG-3′) and *Chdh*-downstream (target sequence: 5′- AGCCGCGCTGTTGCCAGTGTGGG-3′) where underlining indicates the PAM sequences] were *in vitro* synthesized (T7 gRNA Smart Nuclease Synthesis Kit, System Biosciences) following the manufacturer’s instruction. Injected zygotes were transplanted into the uterus of pseudo-pregnant dams, and the targeted region of the *Chdh* gene from the resultant pups, obtained by cesarean section, were examined by direct-sequencing of PCR products amplified from the template DNAs extracted from the tail, with the primer set A (forward: 5′- GCTAGGCATAACCAACCAGC-3′, reverse: 5′-TAGGTCCTGCCTCTAGCAGC-3′). Mutated alleles of the promising founders were further analyzed by sequencing of the PCR products subcloned into pCR2.0 (Invitrogen). When the selected founder (#509) with the desired (loss-of-function) mutation, reached sexual maturity, *in vitro* fertilization was performed with B6N strain-derived oocytes, to obtain mice with heterozygous mutated allele. The heterozygous males and females were intercrossed to produce homozygotes and control littermates. Routine genotyping of mice was done by PCR using the primer set A producing 362 and 319 bp fragments from the wild-type and mutated allele, respectively. To probe a depletion of the Chdh protein in homozygotes, tissues were lysed in the lysis buffer [50 mM Tris-HCl; pH 7.5, 150 mM NaCl, 1% Triton X-100, 0.05% SDS, 1 mM EDTA, 1 mM PMSF and 2 μg/ml aprotinin] by homogenization and sonication, and precleared by centrifugation for 15 min at 4 ºC, followed by SDS-PAGE and Western blotting using anti-Chdh antibody (Thermo Fisher, PA5-50378) and anti-Gapdh antibody (Santa Cruz, sc-20357).

Of the multiple candidate founders obtained, the founder #509 harbored two differently mutated alleles (43-bp and 26-bp deletions downstream of the start codon) in the gene (Supplementary Figure S3A). Both deletions were expected to elicit frameshift of the transcript. Germline transmission of the two deletions was confirmed by mating the founder with inbred B6N, and it was decided to generate and analyze the homozygotes harboring the 43 bp-deletion by mating the heterozygotes. Homozygotes for this mutation were confirmed to lack an expression of the Chdh protein in the kidney where the Chdh protein was abundantly expressed. The expression of Chdh was not detectable with the presently used antibody in the brain from wild-type (WT) (Supplementary Figure S3B), suggesting low level of Chdh expression in the brain. Collectively, the "43bp-deleted allele" was considered to be functionally null.

**Estimation of betaine and other metabolites**

Choline, methionine (Met), betaine, cystathionine, cystine, S-adenosylmethionine (SAM), and S-adenosylhomocysteine (SAH) levels in the brain (frontal cortex), plasma, and kidneys were estimated using Acquity ultra performance liquid chromatography (Waters) coupled to TSQ-Vantage triple-quadrupole tandem mass spectrometry (LC/MS) with multiple reaction monitoring mode (Thermo Fisher).

**Pretreatment for LC-MS analysis**

LC-MS was used to measure following compounds: choline, methionine (Met), betaine, cystathionine, cystine, S-adenosylmethionine (SAM) and S-homocysteine (SAH). To measure betaine and Met, betaine-d11 and methionine-d11 were added to the homogenate or plasma as the internal standards.

***Cortex:*** Tissues (10-20 mg) were homogenized in water by sonication. Three volumes (180 µl of a mixture of chloroform:methanol (1:2) were added to the homogenate, followed by vortexing. Then, 60 µl each of chloroform and water was added for degreasing, followed by vortexing. The mixture was centrifuged (10000 rpm, 4℃) for 15 min. The upper phase (150 µl) was collected, and dried up under $N_2$ gas. After dissolved in 100 µl of 0.1 % formic acid, the sample was filtered through a self-made StageTip C8.

***Plasma:*** Blood samples (0.6-0.8 ml) taken by cardiac puncture were collected to tubes containing 1 mg EDTA, and left for 1 hr at RT. The plasma was collected by centrifugation at 3,400 rpm for 10 min. Three volumes (60 µl) of a mixture of chloroform:methanol (1:2) were added to the plasma, followed by vortexing. Then, 20 µl each of chloroform and water was added for degreasing, followed by vortexing. The mixture was centrifuged (10000 rpm, 4℃) for 15 min. The upper phase (50 µl) was collected was dried up under $N_2$ gas. After dissolved in 100 µl of 0.1 % formic acid, the sample was filtered through StageTip C8.

***Kidney:*** Tissues (50-60 mg) were homogenized in ten volumes of water by sonication. After diluted (1:10) with water, the homogenate was treated by the same procedure as in the brain sample.

The LC condition for the LC-MS analysis using the ACQUITY UPLC (Waters, Milford, MA, USA) is as follows: mobile phase A; 0.1% formic acid; mobile phase B; acetonitrile, flow rate; 0.3 ml/min, gradient; 0/0-4/30-4.1/95-7/95-7.1/0-10/0 (min/B%), column; Triart C18, 2 x 100 mm, YMC Co., Kyoto, Japan), and injection volume; 1 µl. The MS condition is as follows: system; TSQ Vantage EMR (Thermo Fisher), polarity; positive, spray voltage; 3,500 V, vaporizer temperature; 450℃, sheath gas pressure; 50 psi, aux gas pressure–; 15 psi, collision gas pressure; 1 mTorr. Selected-reaction monitoring (SRM) mode was used for quantification. Pre-specified SRM transitions are as follows: choline, m/z 104.2/60.5; betaine, m/z 118.2/42.6 and 58.5; cystathionine, m/z 223.1/88.3 and 134.2; cystine, m/z 241.0/74.3 and 152.1; SAM, m/z 399.1/136.2 and 250.1; methionine, m/z 150.1/56.5 and 104.3; SAH, m/z 385.1/134.1 and 136.2; betaine-d11, m/z 129.1/46.6 and 66.5 and methionine-d11, m/z 153.1/56.5 and 107.3.

**HPLC analysis**

Cysteine (Cys), homocysteine (Hcy) and glutathione (GSH) from the cortex and plasma were measured as their reduced forms by electron chemical detection (ECD-700 HPLC, Eicom, Kyoto, Japan).

***Cortex*****:** Seven microliters of 0.1 M PBS (pH7.2) and 3 µl of 100 mM dithiothreitol (DTT) were added to 50 µl of the homogenates, and then the mixture was placed on ice for 40 min. For deproteinization, 60 µl of 0.4 M perchloric acid was added to the mixture. After placed on ice for 30 min, the mixture was centrifugated (10,000 rpm, 4℃, 15 min). The collected supernatant (100 µl) was diluted with 0.1M phosphate buffer (pH2.5) (1:200), and then subjected to the HPLC analysis.

***Plasma*****:** Nine microliters of 0.1 M PBS (phosphate-buffered saline; pH7.2) and 1 µl of 100 mM DTT were added to 10 µl of the plasma collected, and then the mixture was placed on ice for 40 min. For deproteinization, 20 µl of 0.4 M perchloric acid was added to the mixture. After placed on ice for 30 min, the mixture was centrifugated (10,000 rpm, 4℃, 15 min). The collected supernatant (30 µl) was diluted with 0.1M phosphate buffer (pH 2.5) (1:200), and then subjected to the HPLC analysis.

The HPLC analyses were done by using the EP-700 system (Eicom, Kyoto, Japan) as follows: column; Eicompack SC-5ODS (2.1 mm x 150 mm), detector, ECD-700 (Eicom), mobile phase; 0.1 M phosphate buffer (pH2.5), containing 5mg/l EDTA 2Na, 500mg/l sodium octanesulfonate, and 2.56% (for Hcy) or 1.53% (for GSH and Cys) acetonitrile, flow rate; 0.25 ml/min, injection volume; 5µl, electrode; Gold electrode (WE-Au, Eicom), reference electrode; Ag/AgCl (RE-500, Eicom), applied voltage; 600 mV vs Ag/AgCl and column temperature; 30 ℃.

**Behavioral analysis**

All behavioral tests relevant to psychiatric illnesses, except for the methamphetamine (MAP)-induced behavioral sensitization test, were performed according to the previously published methods [3]. For the MAP-induced behavioral sensitization test, mice were treated without (water) or with betaine (2.17 %) via drinking water for 3 weeks before the first MAP injection (1 mg/kg, s.c., day 1). Betaine supplementation was continued during the experiment period. After the first injection, the MAP injection was repeated once a day for 4 days to induce a sensitization to MAP. Mice were administered the challenge injection of MAP (1 mg/kg, s.c.) at day 15 with a 10-day interval from the fifth injection, and locomotor activity was measured according to the previously published methods [3].

**Methylation analysis in Chdh knockout mouse by targeted bisulfite sequencing**

DNA methylation in frontal cortex of *Chdh* knockout mice ($n$ = 6) and WT control ($n$ = 6) was performed by targeted methylation sequencing. For targeted methylation sequencing (Agilent SureSelectXT Methyl-Seq Target Enrichment kit, Agilent) 109 Mb of mouse genomic regions were enriched, which included CpG islands, known tissue-specific differentially methylated regions (DMR), open regulatory annotations, and Ensembl regulatory features (CpG shores and shelves, DNase I hypersensitive sites, histone modification sites, and transcription factor binding sites). Briefly, DNA was extracted from the frontal cortex of *Chdh* knockout mice ($n$ = 6) and WT control ($n$ = 6) (QIAamp Fast DNA Tissue Kit, Qiagen) as per the manufacturer's protocol. DNA concentration was measured (Qubit 2.0 Fluorometer, Life Technologies), and 1 μg of genomic DNA was fragmented (Covaris S2 ultrasonicator, Covaris Inc.) to yield fragments size of 100–175 bp, which was further end-repaired, 3′ adenylated, and ligated to the methylated adapters. These DNA libraries were hybridized to SureSelect Methyl-Seq capture probes, and the hybrids were captured on streptavidin beads, which were further eluted for bisulfite conversion (EZ DNA Methylation-Gold Kit, Zymo Research). The bisulfite converted libraries were PCR amplified, indexed, and pooled for multiplexing. The sequencing libraries were evaluated (2100 Bioanalyzer, Agilent) and quantified (Qubit 2.0 Fluorometer, Life Technologies), then further sequenced at 100 bp paired end read format on the Illumina sequencing platform (HiSeq2500 platform, Illumina).

The quality of the bisulfite-converted sequencing reads was assessed with FastQC [4]. The adaptor sequences and low-quality bases were trimmed using FASTX tool kit [5], and the trimmed reads were mapped to the mouse genome (mm10) by Bismark software (v.0.10.1) with bowtie (v.1.0.0) using default settings [6]. After de-duplication, the mapped reads were used for extraction of methylated cytosines by the Bismark methylation extractor. Methylation rate was calculated as number of methylated cytosine/ (number of methylated cytosines + number of unmethylated cytosines). Differential methylation between *Chdh* knockout and WT control mice were further tested in the specific regions, such as promoter of the genes, body of the genes, and CpG islands by aggregating the methylation rates at individual sites (only CpG sites) having the read depth (number of methylated cytosines + number of unmethylated cytosines) ≥ 5. Median methylation rate of individual CpG sites was considered as the methylation rate in a defined region (e.g., promoter region). Significant differences in the methylation rate between the *Chdh* knock out and wild type mice were evaluated by two-tailed Student's *t*- test, which was corrected for multiple testing by Benjamini–Hochberg method and a Q-value < 0.05 was considered as statistically significant. A fold change >1.667 and <0.6 was considered as hypermethylated and hypomethylated, respectively. Differentially methylated genes were tested for gene ontology enrichment and pathway analysis. Gene ontology enrichment analysis was performed in PANTHER Overrepresentation Test (http://pantherdb.org/webservices/go/overrep.jsp, annotation version and release date: GO Ontology database; released on 2018-06-01). Reported *P*-values were corrected for multiple testing using the FDR method. Enrichment of pathways was performed using Ingenuity

Pathway Analysis (IPA) (Qiagen, content version: 36601845; release date: 2017-06-22). Statistical significance of the enriched canonical signaling pathways was calculated using Fischer's exact test. A $P$ value < 0.05 was considered as statistically significant.

**RNA-seq analysis**

Total RNA from the frontal cortical brain region was extracted (miRNeasy Mini Kit, Qiagen), and quantity and quality of RNA was estimated (2200 TapeStation system, Agilent). Samples with RNA integrity number (RIN) ≥ 8.7 were further used for library preparation with 200 ng of total RNA as per manufacturer's instructions (TruSeq Stranded mRNA sample prep kit, Illumina). Briefly, poly-A containing mRNA was purified using poly-T oligo attached magnetic beads, which were then heat fragmented and reverse transcribed into first strand cDNA using reverse transcriptase and random primers. Second strand cDNA synthesis was performed by incorporating dUTP followed by the addition of a single "A" nucleotide at the 3′ ends of the blunt fragments to prevent double-stranded cDNA from ligating. After adapter ligation (includes multiplexing barcodes), the cDNA fragments were enriched by PCR (15 cycles) to create the final cDNA library. The quality, size distribution, and quantity of cDNA libraries were assessed (2100 Bioanalyzer, Agilent), which was further sequenced at 100 bp, paired-end read format on the HiSeq2500 platform (Illumina).

The RNA-seq data for the individual samples were de-multiplexed using the unique index adapters. Quality of the sequence reads were evaluated by FastQC (http://wwwbioinformaticsbabrahamacuk/projects/fastqc), and the reads were trimmed for adapter sequences and low-quality bases using FASTX tool kit (http://hannonlabcshledu/fastx_toolkit/indexhtml). The reads were further mapped to the mouse reference genome (GRCm38/mm10, http://hgdownload.soe.ucsc.edu/goldenpath/mm10/chromosomes/) using TopHat with default parameters (v.2.0.14) [7], which utilizes the aligner, Bowtie2 (v.2.2.5) [8]. The expression levels were quantified using Cufflinks (v.2.2.1) [7] based on the read mapping, calculated as fragments per kilo base of transcript per million mapped reads (FPKM), corresponding to the UCSC gene annotations for mm10 (http://hgdownload.soe.ucsc.edu/goldenpath/mm10/database/refFlat.txt.gz). To test the statistical significance for differential expression among the comparison groups, Student's *t*-test on log-transformed FPKM values (log2 FPKM) was applied. A *P*-value < 0.05 was considered as statistically significant and was further analyzed for gene ontology enrichment and pathway analysis. Visualization of differentially expressed genes using volcano plots and principal component analysis of gene expression between different study-groups was done in R (https://www.r-project.org). Differentially expressed genes were tested for gene ontology enrichment and pathway analysis. Gene ontology enrichment analysis was performed in PANTHER Overrepresentation Test (http://pantherdb.org/webservices/go/overrep.jsp, annotation version and release date: GO Ontology database; released on 2018-06-01). Reported *P*-values were corrected for multiple testing using the FDR method. Enrichment of pathways was performed using Ingenuity Pathway Analysis (IPA) (Qiagen, content version: 36601845; release date: 2017-06-22). Statistical significance of the enriched canonical signaling pathways was calculated using Fischer's exact test. A *P* value < 0.05 was considered as statistically significant.

**Real-time quantitative reverse transcription (RT)-PCR**

For targeted gene expression, total RNA was extracted from the mice and human brain samples (miRNeasy Mini Kit, Qiagen), and single stranded cDNA was synthesized using reverse transcription kits (High Capacity RNA-to-cDNA Master Mix, Applied Biosystems and SuperScript VILO Master Mix, Invitrogen). The mRNA levels were determined by real-time quantitative RT-PCR, performed in triplicates, based on the standard curve method. TaqMan probes and primers for the target genes and *GAPDH/Gapdh* (internal control) were selected from TaqMan Gene Expression Assays. Values outside mean ± 2 SD in the group were considered outliers and omitted from the analyses. Significant changes in the target gene expression levels between the test and controls were detected by Student *t*-test (two-tailed) for human brain samples or Bonferroni multiple comparison tests for mouse samples, and $P < 0.05$ were considered as statistically significant.

**Rat primary cortical neuron culture**

Rat primary cortical neuron was isolated from the cortices of Sprague-Dawley rats (obtained from Japan's Charles River) at embryonic day 18.5 (E18.5) as described previously [9] (Supplemental Figure 8B). The cells were placed on poly-D-lysine-coated 96-well plate (for image analysis) or 12-well plate (for RNA isolation) at a density of $1.5\times10^5$ cells/$cm^2$. The neurons were maintained in a serum-free medium [Neurobasal medium (Invitrogen, catalog no: 21103049) supplemented with 0.5 mM glutamine, and B27 supplement (Invitrogen)] until days in vitro (DIV) 7. After DIV 8, media was placed by MEM based media [MEM (Gibco, catalog no: 11090-081) supplemented with 0.5 mM glutamine, and B27 supplement (Invitrogen), 0.06 µM holo-Transferrin (Calbiochem), and 10 mM HEPES (pH7.4)]. The neuron culture was maintained in the presence or absence of PCP (1 µM) and betaine (5, 50, and 500 µM) after DIV 14, and 30 % volume of media were replaced by a new media which contains the identical concentration of PCP and betaine every two days.

**Establishment of GLO1-deficient hiPSCs**

To establish the isogenic *GLO1*-deficient hiPSCs, it was first confirmed that there were no missense variants in the genome of the subject (mentally healthy) for the genes in the carbonyl stress pathway, for example, *GLO1*, *HAGH* (encoding glyoxalase II), *AGER* (encoding advanced glycosylation end-product specific receptor), *LGALS3* (encoding galectin-3), *DDOST* (encoding oligosaccharyl transferase subunit 48), and *PRKCSH* (encoding protein kinase C substrate 80K-H). Human-induced pluripotent cells (hiPSCs) were established from peripheral blood mononuclear cells (PBMCs) using Sendai virus vector as previously described [10]. In brief, PBMCs were isolated from the peripheral blood of healthy control (HC-008) (Leucosep™, Greiner Bio-One). PBMCs were transduced following the manufacturer's instructions (CytoTune®-iPS 2.0 Sendai Reprogramming Kit,

Thermo Fisher). hiPSC culturing was performed as described previously [11, 12]. To generate *GLO1*-deficient iPSCs, a guide RNA sequence (5′-CTTGGTACTGGGGTCCGCGTCGG-3′) was used to target exon 1 of the *GLO1* gene. The gRNA was synthesized using *in vitro* transcription kit (Guide-it™ sgRNA, Takara Bio). The established iPSCs was transfected with 2 µg Cas9 protein (Takara Bio), 0.4 µg gRNA, and 1 µg puromycin-resistant vector (Neon, Thermo Fisher). After puromycin selection, iPSCs were cloned and plated on 96-well plates. After 10 days, single colonies were manually picked and expanded using a culture system (Cellartis® DEF-CS™ 500, Takara Bio). Genomic DNA from single clones was purified using Direct PCR Lysis Reagent (Viagen Biotech). A region containing the target site was amplified and sequenced for mutations in the *GLO1* gene.

**Carbonyl stress assay**

The hiPSCs were cultured in the presence (5, 50 or 500 µM) or absence of betaine for 4 days, then lysed in the lysis buffer (Cell Signaling Technology) with sonication (30 s). Precleared lysates (25 µg protein) were analyzed by Western blotting using anti-AGE (TransGenic), anti-GAPDH (Sigma), or anti-GLO1[13] antibody. Since this anti-AGE antibody has been proven to react mainly to carboxymethyllysine (CML), a subtype of AGEs, this antibody was used as anti-CML antibody. To measure AGEs in the postmortem brains samples, frozen brain samples (BA17) (Fukushima Postmortem Brain Bank, Japan) were lysed in the lysis buffer (Cell Signaling Technology) with sonication (30 s). Precleared lysates (25 µg protein) were analyzed by Western blotting using anti-CEL (TransGenic) and anti-GAPDH (Sigma) antibodies.

**Statistics**

All values in the figures represent the mean ± SEM. Statistical analysis and graphical representation were performed using GraphPad Prism 6 (GraphPad Software). The total sample size (*n*) was described in the respective figure legends. Statistical significance was determined using a two-tailed Student's *t* test. When multiple comparisons were needed, one-way or two-way ANOVA with Fisher's least significant difference (LSD) test, Tukey's multiple comparison test, Bonferroni's correction, or Dunnett's multiple comparison test was used as indicated in the figure legends. A *P* value of less than 0.05 was considered as statistically significant.

### List of Supplementary Tables

**Supplementary Table 1**: Blood biochemistry of *Chdh*-deficient mice

**Supplementary Table 2:** Summary of differentially methylated regions in frontal cortex of *Chdh-deficient* mice

**Supplementary Table 3:** Gene ontology enrichment analysis of differentially methylated gene promoters and gene body ($P < 0.05$) in the frontal cortex of *Chdh*-deficient mice

**Supplementary Table 4:** Summary of differentially expressed genes from the frontal cortex of *Chdh*-deficient mice

**Supplementary Table 5:** Canonical pathways enriched for dysregulated genes in the frontal cortex of *Chdh*-deficient mice

**Supplementary Table 6:** Differentially expressed genes in B6N and C3N mouse strains upon betaine administration

**Supplementary Table 7:** Gene ontology enrichment analysis of genes whose expression positively correlated with principal component analysis (PCA) loading factor in the principal components 3 and 4

**Supplementary Table 8:** Gene ontology enrichment analysis of differentially expressed genes in the frontal cortex of B6N and C3N mouse strains upon betaine administration.

**Supplementary Table 9:** Canonical pathways enriched for differentially expressed genes in the frontal cortex of B6N and C3N mouse strains upon betaine administration.

**Supplementary Table 10:** Comprehensive panel of genes involved in antioxidant activity and the closely coupled, proinflammatory process, tested for mRNA expression levels

**Supplementary Table 11:** Demographic data and sample characteristics of postmortem brain samples (Brodmann Area 17) from individuals with schizophrenia

**Supplementary Table 12:** Correlation between betaine contents and sample characteristics of postmortem brain samples (Brodmann Area 17) from individuals with schizophrenia

**Supplementary Table 13:** Symptom scores of individuals with schizophrenia (postmortem samples) at 3 months prior to the death, rated using the Diagnostic Instrument for Brain Studies (DIBS)
